# 3D-bioprinted skeletal muscle with tailored myofibre architecture

**DOI:** 10.64898/2026.10.18.759932

**Authors:** F. Lezcano, M. Rovira, J. Fuentes, S. Álvarez, S. Sánchez

## Abstract

Engineered skeletal muscle tissue (eSMT) contractile function strongly depends on the alignment and orientation of its myofibres, as force generation and transmission are intrinsically linked to its microstructural architecture. However, fabricating truly functional eSMT with controlled myofibre architecture remains a major challenge. Here, we present a method that combines extrusion-based 3D bioprinting with custom multi-pillar anchoring systems to guide myofibre development. Using this approach, we successfully fabricated self-supporting, functional eSMTs with parallel, fusiform, bipennate, and circular architectures. We then integrated these basic designs into a functional system comprising multiple 3D-bioprinted eSMTs replicating the morphology and myofibre architecture of facial musculature. By recapitulating native muscle architecture, this approach advances the engineering of functional eSMT for multiple biomedical applications and establishes a foundation for next-generation biohybrid robotics.

## Main Text

Skeletal muscle tissue (SMT) is mainly composed of long, multinucleated cells specialized for contraction, known as myofibres (*1*). Its contractile function strongly depends on the alignment and orientation of these myofibres, as force generation and transmission are intrinsically linked to its highly organized, anisotropic architecture (*2*). In engineered SMT (eSMT), recapitulating this structure-function relationship is essential for predictable biomechanical performance. Over the past decade, eSMT has been progressively refined for a broad range of applications, including tissue grafts in regenerative medicine (*3–9*), *in vitro* platforms for drug testing (*10–15*), laboratory-cultivated meat (*16–20*), and biological actuators in biohybrid soft robotics (*21–28*). However, despite significant advances in tissue engineering, fabricating eSMT with tailored myofibre architecture and function remains a major challenge (*29*).

To date, myofibre orientation has mainly been controlled by using substrates or scaffolds containing topographical cues or stiffness anisotropy (*30–33*). In two-dimensional (2D) cultures, controlled myofibre orientation has been achieved by seeding myogenic cells onto patterned surfaces (*30*, *32*, *33*). However, these systems do not fully reproduce the functionality of three-dimensional (3D) tissue, which support longer-term culture, direct assessment of contractile function, and enhanced maturity (*34*). Scaffolds incorporating topographical cues have also been introduced for 3D eSMTs (*31*). Nonetheless, scaffolds provide stiff elements that restrict contraction, and cellular infiltration can result in necrotic regions. Using extrusion 3D bioprinting, a cell-laden hydrogel was 3D-bioprinted onto a supporting mould to fabricate fusiform eSMT with a pennate myofibre angle (*35*). Although promising, myofibre orientation was only partially controlled and contractile performance remained significantly low. More recently, a method for generating anisotropy from droplets was established, enabling the formation of aligned myofibres, although limited to simple architectures (*36*).

3D bioprinting enables the precise arrangement of cells and biomaterials within a pre-designed structure, allowing the fabrication of reproducible and scalable tissues (*37*). Extrusion-based 3D bioprinting of myogenic cell-laden bioinks promote cell alignment along the print path, providing a potential strategy to pre-define myofibre architecture (*38*). However, controlling print path directionality becomes increasingly difficult as model complexity increases. To address this challenge, new programming tools now allow users to define every segment of the print path and all its printing parameters (*39*). Additionally, an alternative to topographical cues to guide myofibre orientation during myofibre development is to apply mechanical stress along the desired direction of alignment (*40–42*). By generating mechanical stress within the tissue, the anchoring of eSMT promotes cell orientation and myogenic development while also preventing tissue collapse (*3*). In native tissue, SMT is anchored to the skeleton through tendons, which exhibit different architectures depending on the shape and function of the muscle (*43*). However, artificial anchoring systems described in the literature typically rely on relatively simple designs, usually consisting of pairs of rectangular or cylindrical pillars, therefore supporting basic eSMT configurations, mainly strips or rings of tissue (*3–6*, *10–14*, *21–26*, *36*, *44–46*). To support more complex tissue architectures, such as parallel, fusiform, pennate or circular configurations, more complex anchoring systems are required.

We hypothesized that combining precise control over the extrusion-based print path with custom multi-pillar anchoring systems to guide myofibre development along predefined directions would enable the fabrication of complex eSMTs with tailored myofibre architectures. Based on this hypothesis, we developed a new method, through which we successfully fabricated self-supporting, biomimetic eSMTs with parallel, fusiform, bipennate, and circular architectures for the first time. We then combined these basic designs to obtain a functional system composed of multiple 3D-bioprinted eSMTs that replicated the morphology and myofibre architecture of facial musculature (Figure S1). By recapitulating native muscle architecture, this approach advances the fabrication of functional eSMT for multiple biomedical applications and sets the stage for next-generation biohybrid robotics.

### Effect of anchoring system design and print path direction on myofibre orientation

We tested different anchoring systems and print paths to investigate how they affected eSMT myofibre development. We designed two different anchoring systems, one consisting of two standard rectangular pillars and the other consisting of six thin cylindrical pillars (Fig. 1A). We then designed rectangular eSMTs for bioprinting around these anchoring systems following different print paths, one aligned with the direction of the desired muscle fibres (axial direction) and the other perpendicular to it (transversal direction) (Fig. 1A). The thick black arrows in Figure 1A indicate the direction of the stress induced by each anchoring system within the tissue, showing that the six thin cylindrical pillars produce stress along more parallel directions.

**Fig. 1.**
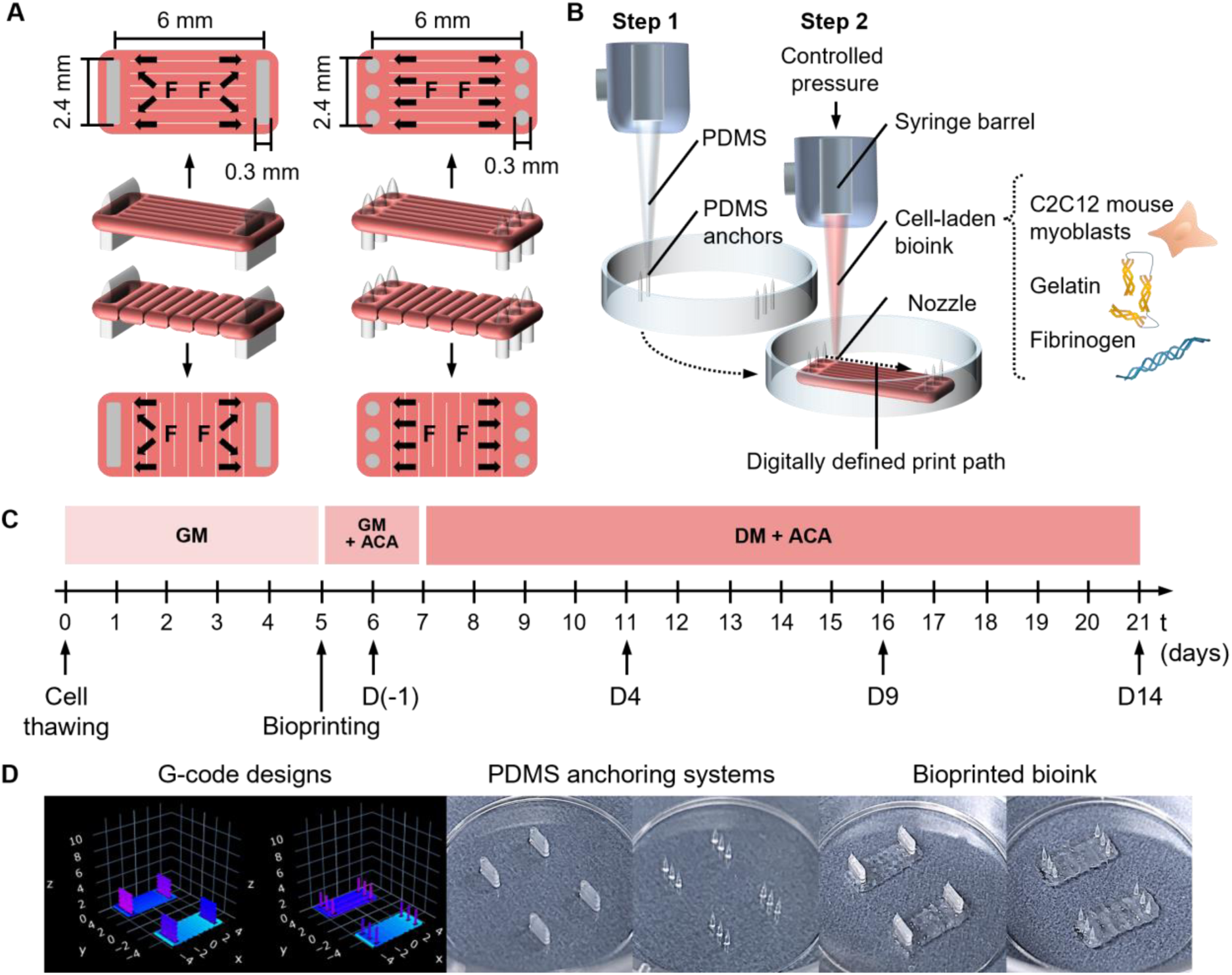
Design and fabrication of anchoring systems and eSMT. (**A**) Schematics of the different anchoring systems, print path direction, and mechanical stress direction within the tissue (F; thick black arrows). (**B**) Schematics of the extrusion-based 3D printing process and bioink composition. (**C**) Tissue biofabrication and maturation timeline, where D(−1), D4, D9, and D14 are the characterization timepoints. (**D**) Representation of the print paths defined as G-codes, 3D printed PDMS anchoring systems, and bioprinted cell-laden bioink around previously cured anchors.

We fabricated all anchoring systems and eSMTs using an extrusion-based bioprinter (Fig. 1B). In a first step, we 3D-printed the anchoring systems in PDMS and cured them overnight. In a second step, we 3D-printed the eSMT around the pre-cured anchoring system using a cell-laden bioink composed of C2C12 mouse myoblasts, gelatine, and fibrinogen according to Fuentes *et al.* (*47*). We assessed tissue maturation and myofibre orientation at different time points: 1 day before starting differentiation (D(−1)) and 4, 9, and 14 days after starting differentiation (D4, D9, and D14), as shown in Figure 1C. Figure 1D shows the digitally defined print paths implemented to print the anchoring systems and the eSMTs, which were generated as G-code using a custom-made Python script, which allowed precise control over 3D geometry, print path direction and speed, line width, and layer height (*39*).

Figure 2 shows the evolution of tissue maturation, cell viability, and myofibre orientation over time for the four conditions introduced in Figure 1A: two pillars with axial print path (2PA), two pillars with transversal print path (2PT), six pillars with axial print path (6PA), and six pillars with transversal print path (6PT).

**Fig. 2.**
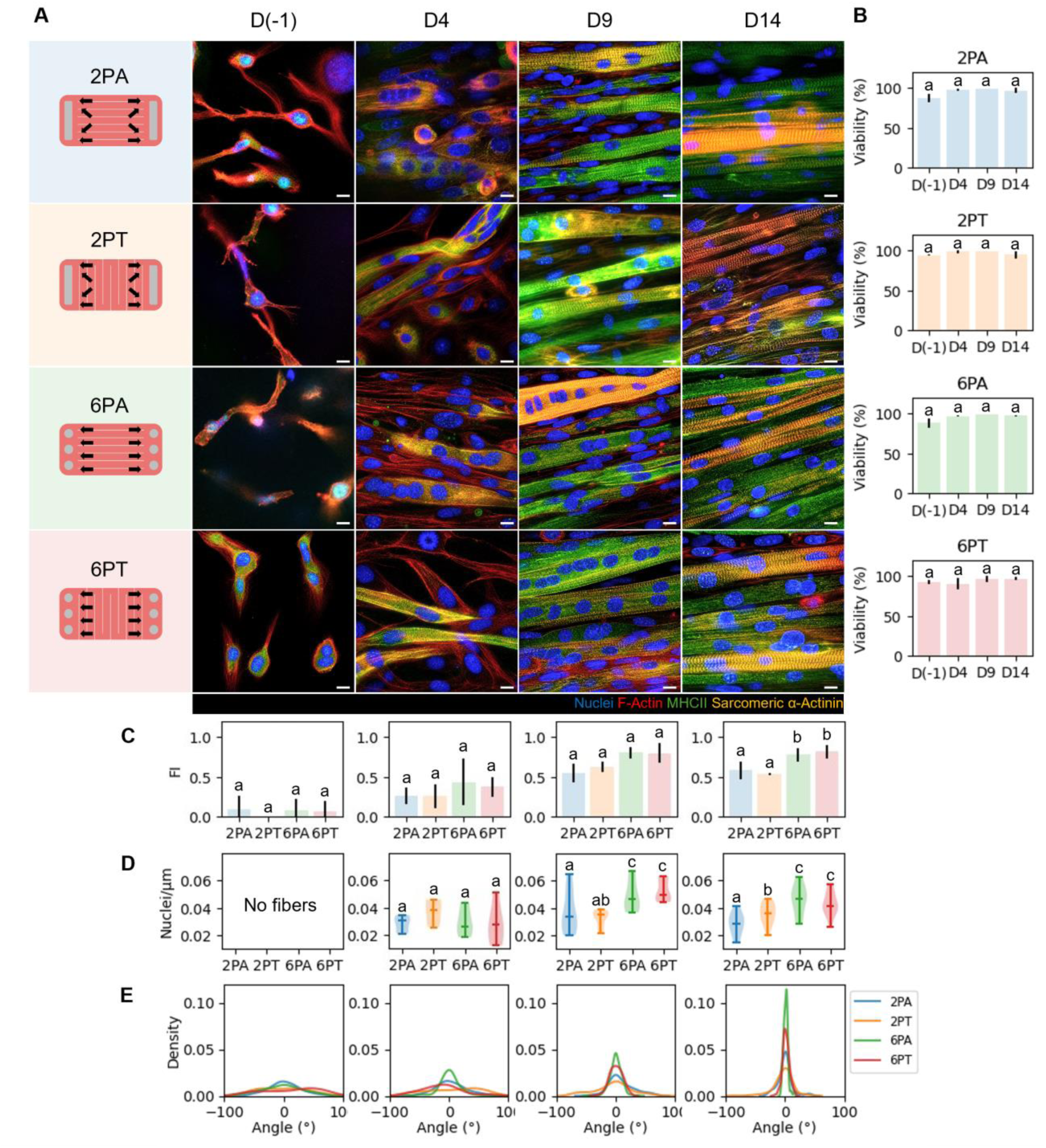
Tissue maturation, cell viability and myofibre orientation over time for different anchoring systems and print path directions. (**A**) Representative fluorescence images (100X) showing cell nuclei (blue), F-actin (red), MHCII (green), and sarcomeric α-actinin (yellow). Scale bar: 10 µm. (**B**) Cell viability. (**C**) Fusion index. (**D**) Nuclei density. (**E**) Kernel density plots of fibre orientation angles over time, where 0° represents the axial direction.

We observed myofibres with well-defined sarcomeres from D9 onwards, with fibre diameter becoming more homogeneous by D14 (Fig. 2A). All conditions showed high cell viability, with values of approximately 90% at D(−1) that increase to almost 100% by D4 (Fig. 2B, Data S1).

However, from D9 onwards, both the fusion index (FI) and nuclei density were higher in all tissues anchored to the six-pillar anchoring systems, suggesting that the thin, cylindrical pillars promote muscle fibre maturation more efficiently (Figure 2C-D, Data S1). At D(−1), FI values reflected the presence of early-stage fibres containing two or three nuclei. However, nuclei density could not be reliably calculated at this time point due to the low number of nuclei per fibre.

We processed fluorescence images of F-actin fibres to assess the evolution of muscle fibre orientation over time. Each vector in Figure S2 represents the orientation of local F-actin filaments, which matches that of the myofibres. Figure 2E shows kernel density plots representing the probability density of the orientation of these vectors for each tissue condition, where 0 degrees corresponds to the axial direction (Data S4). Up to D4, myofibre orientation was mainly influenced by the print path direction, as the kernel density plots showed one predominant direction of alignment for the axial print path and multiple directions for the transverse print path, with clear tendencies towards 90 and −90 degrees. From D9 onwards, the influence of the anchoring system is clearly observed, as all myofibres start aligning along the direction of the mechanical stress produced by the anchors. By D14, the resulting myofibre orientations in all conditions were consistent with the direction of the tension generated within the tissue over the differentiation period. Additionally, the tissues bioprinted with axial print paths showed better alignment than the ones printed with transversal print paths, as indicated by higher and narrower peaks in the density plots.

### Bioinspired design and fabrication of functional eSMT with tailored myofibre architecture

We fabricated eSMTs with parallel, fusiform, bipennate and circular architectures (Fig. 3A) by implementing anchoring systems with thin, cylindrical pillars and print paths in the direction of the desired myofibres (Fig. 3B). We positioned the anchoring pillars in strategic positions to generate mechanical stress in the direction of the desired myofibres, as indicated by the thick black arrows representing the stress in Figure 3B. The implemented print paths, the obtained PDMS anchoring systems, and the eSMTs printed around them are shown in Figure S3.

**Fig. 3.**
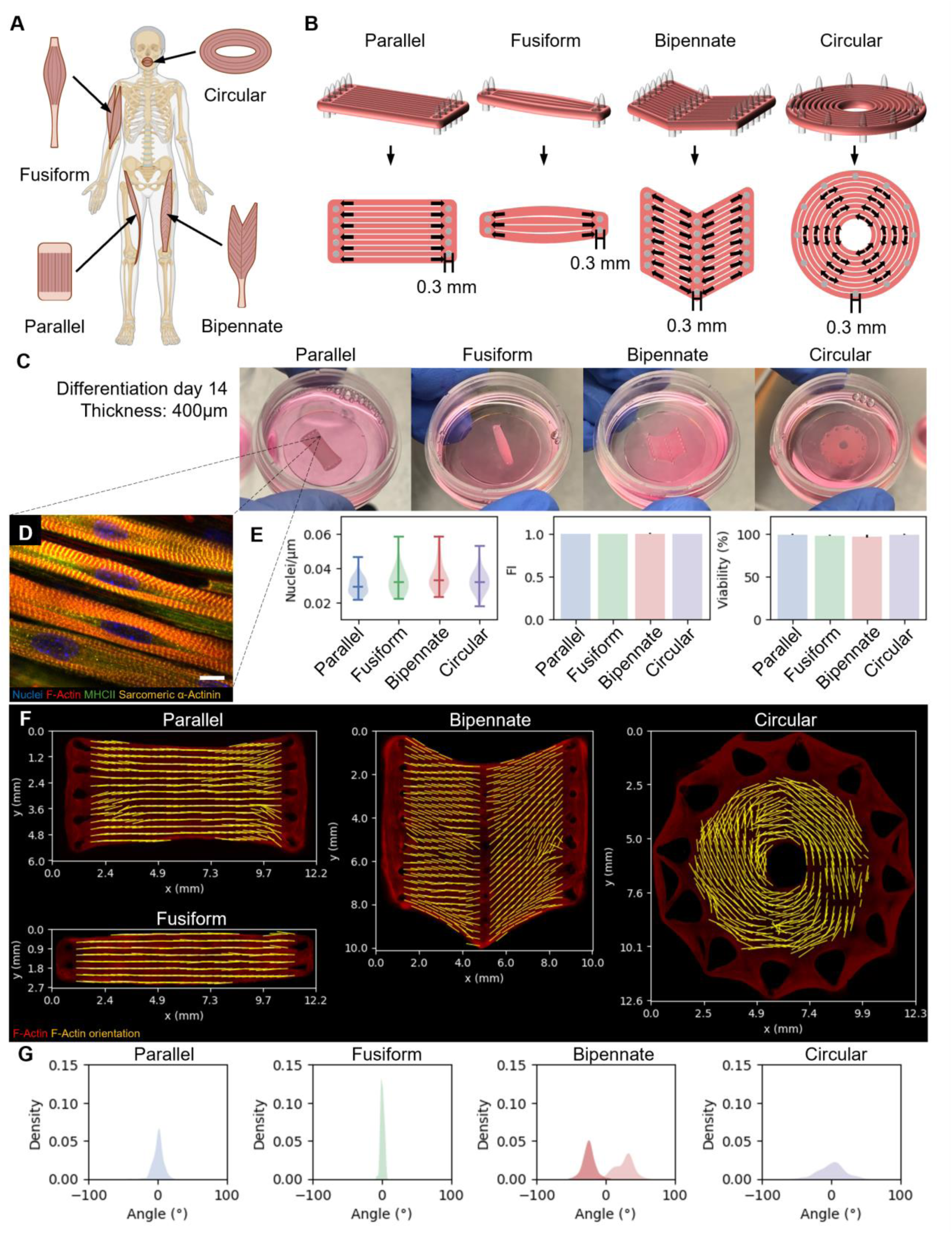
Bioinspired eSMTs design and characterization. (**A**) Basic human skeletal muscle architectures. (**B**) Skeletal muscle construct and anchoring system design, and mechanical stress direction within the tissue (thick black arrows). (**C**) Pictures of the eSMTs in DM after 14 days of differentiation. (**D**) Representative confocal image of myotube structure, showing nuclei, F-actin, MHCII, and sarcomeric α-actinin after 14 days of differentiation. Scale bar = 10 µm. (**E**) Nuclei density, fusion index, and cell viability (n = 3). (**F**) Representative plots of vector fields showing F-actin orientation on top of florescence images (10X) of F-actin fibres (red). (**G**) Kernel density plots of fibre angles, where 0° corresponds to the x direction from panel D, except for the circular design, where 0° represents the direction perpendicular to the radius of the tissue (n = 3). Schematic in panel A was created with BioRender.com.

We cultured all the bioinspired eSMTs in differentiation medium over 14 days (Figure 3C). Over this period, all eSMTs shrank, to approximately half the initial size (Figure S4). Tissue thickness decreased from 800 µm to 400 µm. After the differentiation period, confocal images revealed mature myofibres with elongated nuclei and well-defined sarcomeres (Fig. 3D). Cell viability and nuclei density showed high values, and fusion index showed values within the range reported for native tissue (Fig. 3E, Data S2) (*48*, *49*).

Figures 3F-G show vector fields overlaid on fluorescence images of F-actin filaments (Fig. S5-8), indicating the local orientation of myofibres, and kernel density plots of the orientation of these vectors, respectively (Data S4). This analysis revealed that the myofibres adopted the predefined orientation in all cases, with axial myofibres in the parallel and fusiform configurations, two main myofibre orientations in the bipennate configuration, and myofibres perpendicular to the radial direction in the circular configuration. Parallel and fusiform architectures showed highly aligned fibres, whereas bipennate and circular architectures were less aligned, possibly due to weaker mechanical stress resulting from interactions among multiple pillars.

We also characterized the contractile function of the biomimetic eSMTs at D14. We electrically stimulated all tissues using electrical pulses of different voltage intensities and frequencies (Fig. 4A), using two graphite electrodes placed inside the Petri dish containing the self-supporting tissue detached from its anchors and submerged in culture medium (Fig. 4B). We quantified contraction length by measuring the percentage contraction of the tissue along different directions depending on the tissue design (Fig. 4C). Stimulation at a pulse frequency of 1 Hz induced twitch contractions, while a pulse frequency of 30 Hz induced tetanic contractions (Fig. 4D, Data S3).

**Fig. 4.**
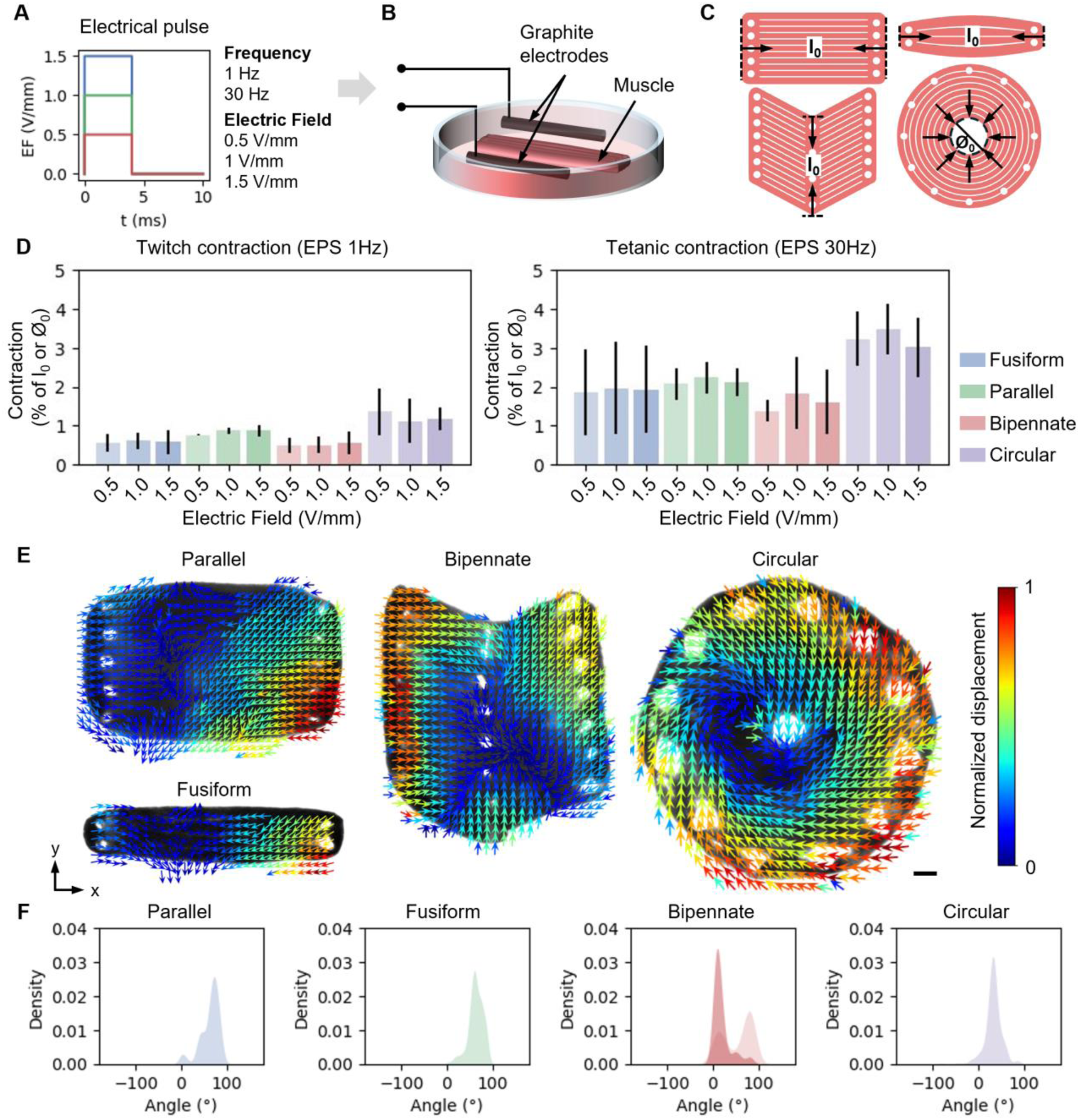
Bioinspired eSMTs contractile function assessment. (**A**) Electrical pulse for twitch (1 Hz) and tetanic (30 Hz) stimulation, with different intensities (red, green, and blue for 0.5, 1, and 1.5 V/mm, respectively). (**B**) Stimulation setup. (**C**) Measurements used to assess contraction, where l0 is the initial length of the parallel, fusiform, and bipennate architectures, and Ø0 is the initial diameter of the circular architecture. (**D**) Muscle contraction as a percentage of l0 or Ø0 for twitch and tetanic stimulation applying different electric field intensities (n = 3). (**E**) Vector field showing the direction of contraction and the normalized displacement with values between 0 and 1, scale bar: 1 mm. (**F**) Kernel density plots of contraction direction angles, where 0° corresponds to the x direction, except for the circular design, where 0° represents the direction perpendicular to the radius of the tissue (n = 3).

We post-processed bright field videos of eSMT contraction (Movie S1) to obtain vector fields showing the local displacement of the tissue (Fig. 4E). Probability density plots of the orientation of these vectors revealed that the direction of contraction (Fig. 4F) and the orientation of the fibres (Fig. 3G) in the tissues presented similar patterns, as the tissues contract according to the orientation of their myofibres (Data S4).

### Assembly of multiple eSMTs with tailored myofibre architecture

We designed and fabricated eSMTs inspired by twelve superficial muscles of the face (Fig. 5A) together with their corresponding anchoring systems following the same approach as in the previous section (Fig. S9) but using a different bioink, incorporating Matrigel and a higher cell concentration (20×10^6^ cells/ml). We cultured all tissues individually in differentiation medium for 14 days. During this period, the muscles matured and compacted acquiring their final shape (Fig. S10). Figure 5B shows all matured eSMTs placed on top of a 3D-printed skull model, depicting their spatial relationship with the underlying skeletal structures. All matured eSMTs presented the predefined, biomimetic myofibre architecture (Fig. S11-14, Fig. 5C). The probability density of myofibre orientation confirmed one main direction for frontalis, procerus, risorius, and zygomaticus major, two main directions for the right-left transverse nasalis system, and a circumferential direction for the orbicularis (Fig. S15).

**Fig. 5.**
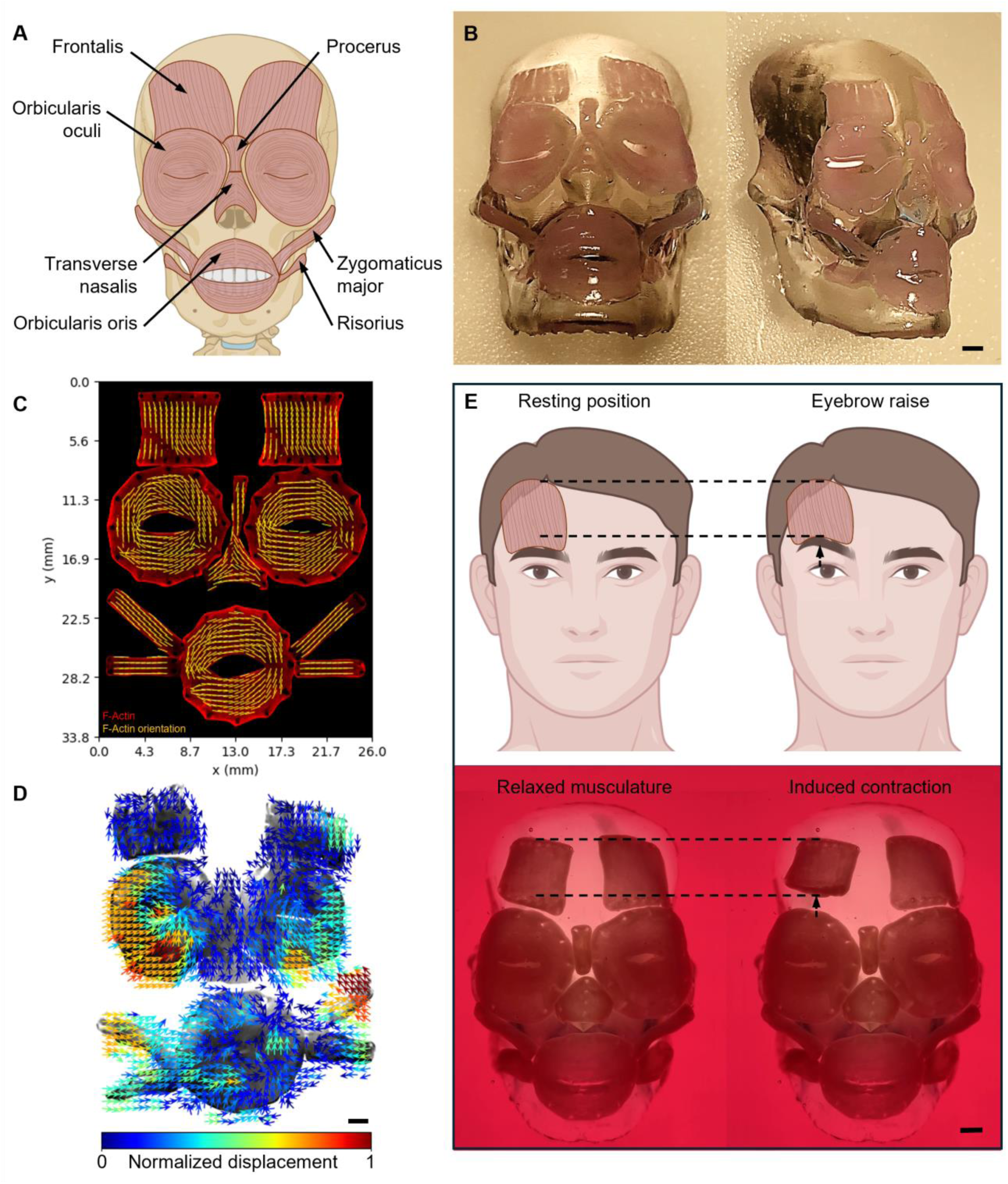
Bioprinted face musculature. (**A**) Twelve superficial muscles of the face. (**B**) Pictures of bioprinted facial eSMTs on top of a 3D printed skull after 14 days in differentiation medium. Scale bar: 2 mm. (**C**) Vector field (yellow) showing F-actin orientation on top of a fluorescence image (10X) of F-actin fibres within the eSMTs (red). (**D**) Vector field showing the local direction of contraction and the normalized displacement of a 2D arrangement of facial eSMTs under tetanic electrical pulse stimulation. Scale bar: 2 mm. (**E**) Schematics of a relaxed face (top left) and a face with the right eyebrow raised (top right), a gesture produced by contraction of the frontalis muscle. Images of relaxed facial eSMTs fixed onto a 3D-printed skull and immersed in culture medium (bottom left), and electrically induced contraction of the right frontalis muscle after detachment of its lower end (bottom right). Black arrows indicate muscle displacement. Scale bar: 2 mm. Schematics in panels A and E were created with BioRender.com.

To analyse the contractile function of the eSMTs, we arranged the self-supporting tissues in the same 2D configuration shown in Figure 5C and placed them inside a petri dish with electrodes as depicted in Figure 4B. Under tetanic electrical pulse stimulation, all eSMTs contracted according to their specific myofibre architecture (Fig. 5D, Movie S2). We used the same stimulation setup to observe tissue contraction in a more natural 3D configuration. We coated a bioprinted skull with a thin layer of fibrin, and placed the tissues on top of it, in their respective anatomical positions as shown in Figure 5B and E. To keep the muscles in their position and prevent them from floating, we fixed them to the surface of the skull using small amounts of fibrin. Although this anchoring strategy significantly restricted muscle contraction, under tetanic electrical stimulation the tissues still contracted in accordance with their specific myofibre architecture (Movie S3). This does not fully reproduce the *in vivo* behaviour of the musculature, as these muscles are normally attached at specific points to soft tissue and to the skin. In native tissue, the inferior fibres of the frontalis insert into the skin and subcutaneous tissue of the eyebrow region. When properly connected to the skin, contraction of the muscle elevates the eyebrow, as shown in the schematics in Figure 5E. We reproduced this by inducing the contraction of the right frontalis eSMT after detachment of its lower end from the skull model. Movie S4 and its frames in Figure 5E show the contractile response of the eSMT to electrical stimulation and its potential to actuate the overlying facial skin, thereby potentially contributing to the reproduction of facial expressions.

## Discussion and conclusion

This work presents a new strategy for engineering functional 3D eSMTs with tailored myofibre architectures by combining multi-pillar anchoring systems with controlled extrusion print paths. Using this approach, we successfully fabricated self-supporting eSMT reproducing the basic architectures and functions of human skeletal muscles for the first time. We subsequently combined these eSMTs into a functional system replicating the morphology, myofibre architecture and contractile function of human facial musculature, demonstrating that the proposed strategy can produce integrated muscular systems with complex geometries and predictable myofibre and contraction patterns.

We show that myofibre development is initially influenced by extrusion print path direction and subsequently by anchor-induced mechanical stress, establishing the basis for a strategy that overcomes the limitations of using substrates or scaffolds with topographical cues (*30–32*). This new strategy enables the development of complex anchoring systems, capable of supporting diverse tissue architectures, such as the parallel, fusiform, bipennate or circular configurations presented here. This represents an advance beyond conventional anchoring systems, which typically consist of pairs of rectangular or cylindrical pillars, supporting simple eSMT configurations, mainly strips or rings of tissue (*3–6*, *10–14*, *21–26*, *36*, *44–46*). Through our approach, we achieved mature myofibres with nuclei density and fusion index values within the ranges reported for native tissue (*48*, *49*). Contractility values substantially exceeded those of current eSMTs with multiple myofibre orientations (*31*, *35*) and fell within the range reported for native tissue (*50*).

This work enables a shift from functionality-driven approaches that overlook tissue architecture towards design-driven strategies that enable functionality through precise architectural control. By recapitulating native muscle architecture, our approach advances the engineering of functional skeletal muscle bioactuators for a wide range of biomedical applications and lays the foundation for next-generation biohybrid robotics.

## Acknowledgments

Funding: The research leading to these results received funding from: MICIU/AEI/10.13039/501100011033 “Centro de Excelencia Severo Ochoa” grant CEX2023-001282-S.

The CERCA program by the Generalitat de Catalunya.

MICIU/AEI/10.13039/501100011033 and ESF+ grant JDC2024-053912-I (FL).

The BIST Ignite program (FL).

Secretaria d’Universitats i Recerca del Departament d’Empresa i Coneixement de la Generalitat de Catalunya through the project 2021 SGR 01606 (SS).

The European Union’s Horizon Europe research and innovation programme under grant agreement No. 101070328 (SS).

The “Constantes y Vitales” 2023 prize (SS).

## Author contributions

Conceptualization: FL

Methodology: FL, JF

Validation: FL, MR

Investigation: FL, MR, JF, SA

Visualization: FL

Software: FL

Funding acquisition: FL, SS

Project administration: FL, SS

Supervision: FL, SS

Writing – original draft: FL

Writing – review and editing: FL, SS

## Data and materials availability

All data needed to evaluate the conclusions in the paper are present in the paper and/or the Supplementary Materials.

## Supplementary Materials

## Materials and Methods

### Design

All anchoring systems and eSMTs were designed using the FullControl package for Python (*51*). Printing height, width, direction and speed were explicitly defined at each point along the print path in G-codes, obtained by implementing custom Python scripts.

Vertical print paths were defined to obtain anchoring systems composed of cylindrical pillars measuring 0.2 mm in thickness and 2 mm in height. The printing speed was set to 25 mm/s for all anchoring systems.

Each eSMT was designed to follow a print path around the pillars and along the direction of the desired myofibres, except for the rectangular eSMTs shown in Figure 1A, for which the print path was perpendicular to this direction. All print path lines were defined with a height and thickness of 0.3 mm, except for the print path used to obtain the fusiform eSMT, which incorporated different line thicknesses to generate a thicker section in the middle of the tissue than at the sides. The printing speed was set to 80 mm/s for all eSMTs, except for the fusiform eSMT, for which slower speeds were used for the thicker print-line segments.

### Anchoring system 3D printing

All anchoring systems were 3D printed in petri dishes (35 mm diameter, glass bottom) with PDMS 10:1 (DOWSIL-SE1700) using an extrusion-based 3D printer (Inkredible+, Cellink AB) and cured at 65°C overnight. Printing pressure was set to 200 kPa and the speed followed the values programmed in the G-code.

### Cell culture and bioink formulations

The bioink used to develop the method and to print the parallel, fusiform, bipennate, and circular bioinspired designs, was formulated according to Fuentes *et al.* (*52*). Cryopreserved C2C12 cells were thawed, seeded in T-175 Flasks (5×10^5^Cells/Flask), and cultured in GM (DMEM, 10% FBS, 1% Penicillin/Streptomycin, 1% L-glutamine) inside an incubator at 37°C and 5%CO_2_ for 4 days. A 70mg/mL stock solution of gelatine in GM with 2mg/mL aminocaproic acid, and a 40mg/mL stock solution of fibrinogen in PBS 1X were prepared and then mixed at a 1:1 rate. The cells were trypsinized, centrifuged and added to the gelatine-fibrinogen solution at a concentration of 5×10^6^ cells/ml. The resulting cell-laden bioink was kept in the fridge at 4 °C for 10 minutes before starting printing.

The face muscles were bioprinted using a different bioink formulation. The bioink was prepared following the procedure described above, with some modifications: the C2C12 cell density was increased to 20×10⁶ cells/mL, while the gelatine and fibrinogen concentrations were reduced to 50 mg/mL and 20 mg/mL, respectively. In an additional step, Matrigel was added to the gelatine/fibrinogen/C2C12 mixture to obtain a final bioink formulation containing 30% Matrigel.

### eSMT 3D bioprinting

All eSMTs were 3D printed in a petri dish (35 mm diameter, glass bottom) around the pre-cured PDMS anchoring systems using an extrusion-based 3D bioprinter (Inkredible+, Cellink AB). The printing pressure was set to 40 kPa. After printing, 500uL of a 1:10 solution of thrombin in PBS 1X was added to the eSMT to initiate the fibrinogen crosslinking. After 10 minutes, the thrombin was replaced with GM, and the bioprinted eSMT was stored inside the incubator at 37 °C and 5% CO_2_.

### eSMT maturation

ll bioprinted eSMTs were cultured for 2 days in GM with 2mg/mL ACA, and afterwards, in DM (DMEM, 10% HBS, 1% Penicillin/Streptomycin, 1% L-glutamine, 0.1% Insulin-like growth factor, 2mg/mL ACA) inside the incubator at 37°C and 5% CO_2_. DM was replaced with fresh DM every 2 days.

### Cell viability assessment

Cell viability was obtained by the computational processing of fluorescence images randomly taken at different tissue depths, covering a total surface of 5% of the tissue size. The nuclei were stained with Hoechst (blue channel), dead cell nuclei were stained with Ethidium Homodimer-1 (red channel), and live cells were stained with Calcein (green channel) following the standard protocol by the manufacturer (LIVE/DEAD Viability/Cytotoxicity Kit for Mammalian Cells, ThermoFisher Scientific). Cellpose (*53*) was used to segment all nuclei in the blue channel (Fig. S15). The number of segmented nuclei defined the total number of cells present. The number of nuclei inside red regions defined the number of dead cells (N_dead cells_). The number of nuclei that were not inside red regions, but were inside green regions, defined the number of viable cells (N_viable cells_). Cell viability (CV) was then calculated as follows:

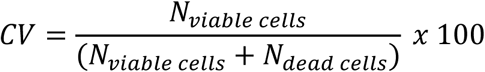

Three eSMTs of each design were analysed.

### Staining

The 3D-bioprinted tissue constructs were washed three times with PBS and then fixed with formalin 10% for 45 minutes at room temperature (RT). The constructs were then washed again three times with PBS and stored at 4°C until further use. For immunostaining, the tissues were permeabilized with 0.3% Triton-X-100 in PBS for 10 minutes at RT. After three washes with PBS, the cells were blocked with 5% of Bovine Serum Albumin (BSA) in PBS (PBS-BSA) for 40 minutes at RT to prevent non-specific binding. The tissues were then washed three times with PBS. A solution with primary antibody Rabbit polyclonal to Sarcomeric Alpha (Abcam; 1:200) and monoclonal antibody Anti-Myosin Heavy Chain Alexa Fluor® 488 (eBioscience; 1:100) in PBS-BSA was added into each sample and incubated overnight at 4°C in dark conditions. After incubation the skeletal muscle constructs were washed three times with PBS. For the second incubation, a solution with secondary antibody Rabbit anti-sarcomeric Alpha Actinin Alexa Fluor® 647 (Invitrogen; 1:300) and ActinRed™ 555 ReadyProbes™ (Invitrogen; 2 drops/mL) in PBS-BSA was added to the samples for 2 hours at RT in dark conditions. After incubation, unbound antibodies were removed by washing three times with PBS. Nuclei were stained with Hoechst 33342 (Life Technologies, 0.5 µg/mL) for 10 minutes. Finally, the samples were washed three times with PBS and stored at 4 °C until imaging. Fluorescently labelled tissue constructs were imaged using an inverted microscope (DMi8, Leica) and a Zeiss LSM 800 confocal scanning laser microscope (CSLM). Excitation wavelengths were set as follows: 488 nm for Myosin Heavy Chain, 405 nm for nuclei, 555 nm for F-actin, and 647 nm for sarcomeric α-actinin.

### Myofibre orientation

Full-tissue fluorescence images (10X) of stained F-actin filaments were analyzed using OrientationJ plugin for ImageJ to obtain a vector field representing the direction of the myofibres. The obtained data was postprocessed using a custom-made python script to obtain kernel density plots with the probability density of vector orientations. Three eSMTs of each design were analysed.

### Fusion index assessment

Fluorescence images (100X) of stained F-actin fibres (red channel), MHCII (green channel), sarcomeric α-actinin, and nuclei (blue channel) were postprocessed using Cellpose and ImageJ. The nuclei were segmented using Cellpose, obtaining the total number of nuclei in the image (N_nuclei_) and a region of interest (ROI) for each nucleus with the sole purpose of individualizing each nucleus. Then, the number of nuclei inside regions occupied by myofibres (N_nuclei inside myofibres_) was manually counted (*54*). The FI was then calculated using the following equation:

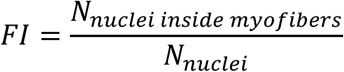

For each design, three images were analysed from each of three independent eSMTs.

### Nuclei density assessment

Fluorescence images (20X) of stained F-actin filaments and nuclei were postprocessed using ImageJ. The length (L) of a fibre segment containing 5 consecutive nuclei was measured as shown in Figure S16 and the nuclei density (ND) was calculated as follows:

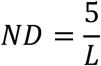

Three eSMTs of each design were analysed, considering 10 fibre segments from each individual eSMT. For parallel, fusiform, bipennate, and circular architectures, 30 fibre segments were considered.

### Electrical pulse stimulation and contractility assessment

Parallel, fusiform, bipennate, and circular eSMTs were electrically stimulated using graphite electrodes according to Mestre *et al*. (*54*) (Figure 5A-B). Each eSMT was transferred to an anchor-free petri dish (35 mm diameter, glass bottom), and the electrodes were submerged into the DM, one at each side of the tissue (Figure 5B). Electric fields with a pulse width of 4 ms and intensities of 0.5 V/mm, 1 V/mm, and 1.5 V/mm were applied to the DM to induce contraction in the eSMT. A frequency of 1 Hz was implemented to induce twitch contractions, and a frequency of 30 Hz was implemented to induce tetanic contractions. An inverted microscope (DMi8, Leica) in bright-field mode (1.25X) was used to record 25-FPS videos of the eSMTs contracting. Twitch contractions were recorded over 10 seconds. Tetanic contraction recordings started with 5 seconds of the eSMT in resting state, followed by 5 seconds of tetanic stimulation and 5 seconds of relaxation without stimulating.

Contractility was assessed by measuring the length of the tissue before (*L*_0_) and during contraction induction (*L*) using ImageJ. The percentage of contraction was calculated using the following equation:

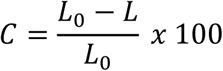

For the circular tissue, instead of measuring the length L, we measured the diameter as shown in Figure 5C. Videos of three eSMTs of each design were analysed.

For electrical stimulation of the facial musculature in 2D and 3D, the eSMT composition was placed in a glass Petri dish containing DM. Individual eSMTs were attached to each other and/or to the 3D-printed skull by locally applying fibrinogen (10 mg/mL), followed by thrombin (5 U/mL), using a micropipette. The assembled musculature was electrically stimulated by placing two graphite electrodes in the DM. An electric field with a pulse width of 4 ms and an intensity of 1.5 V/mm was applied to induce tetanic contraction of the eSMTs. Tetanic contractions were videorecorded at 24 FPS using a smartphone camera.

### Particle Imaging Velocimetry (PIV)

Tetanic contraction videos were postprocessed using PIV plugin for ImageJ in order to obtain a vector field representing the displacement of the eSMT pixels during contraction. The vector field data was postprocessed using a custom-made python script to obtain kernel density plots with the probability density of vector orientations. Three eSMTs of each design were analysed.

### Skull 3D printing

An open-source 3D skull model (3DFactoryMX, 2015) was imported into Autodesk Fusion 360, where minor geometric modifications were performed: the occipital was cut to obtain a flatten surface, and the nose cartilage was added. The modified model was subsequently prepared for additive manufacturing and fabricated using a Form 4B stereolithography 3D printer (Formlabs, USA) with Clear Resin V5 (Formlabs). The skull was printed at a layer height of 0.025 mm to achieve high geometric fidelity and resolution. Following printing, the model was removed from the build platform and post-processed according to the manufacturer’s recommended procedures.

### Skull coating and muscle placement

A fibrinogen (10 mg/mL) and gelatine (50 mg/mL) solution was prepared by separately dissolving fibrinogen (20 mg/mL) in PBS 1X at 4 °C and gelatine (100 mg/mL) in PBS 1X at 42 °C, followed by mixing the two solutions at a 1:1 ratio. In parallel, a thrombin solution (5 U/mL) was prepared in PBS 1X and maintained at 4 °C. The skull was cooled to 4 °C and then submerged in the fibrinogen/gelatine solution. It was subsequently kept at 4 °C for 5 min before being submerged in the thrombin solution to crosslink the fibrinogen. This process was repeated three times. After coating the skull, the muscles were manually placed on its surface and glued at their insertion sites by locally adding fibrinogen (10mg/ml) followed by thrombin (5 U/ml), using a micropipette.

### Statistical analysis

The statistical analysis was carried out using the Python, implementing functions of Matplotlib, Seaborn, Statistics and Scipy libraries. All error bars represent SDs. A Kruskal-Wallis test was applied to identify differences among three or more groups, followed by pairwise comparisons using Dunn’s post hoc test with Bonferroni correction.

**Fig. S1.**
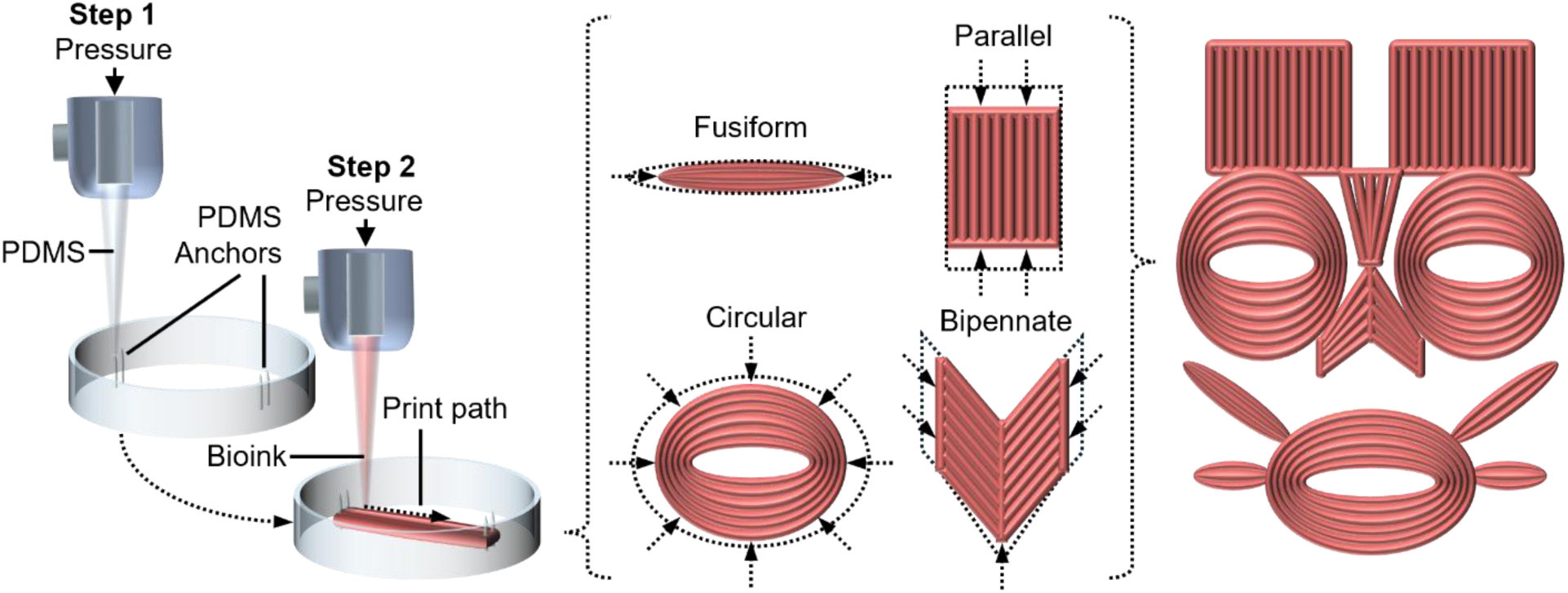
Graphical abstract.

**Fig. S2.**
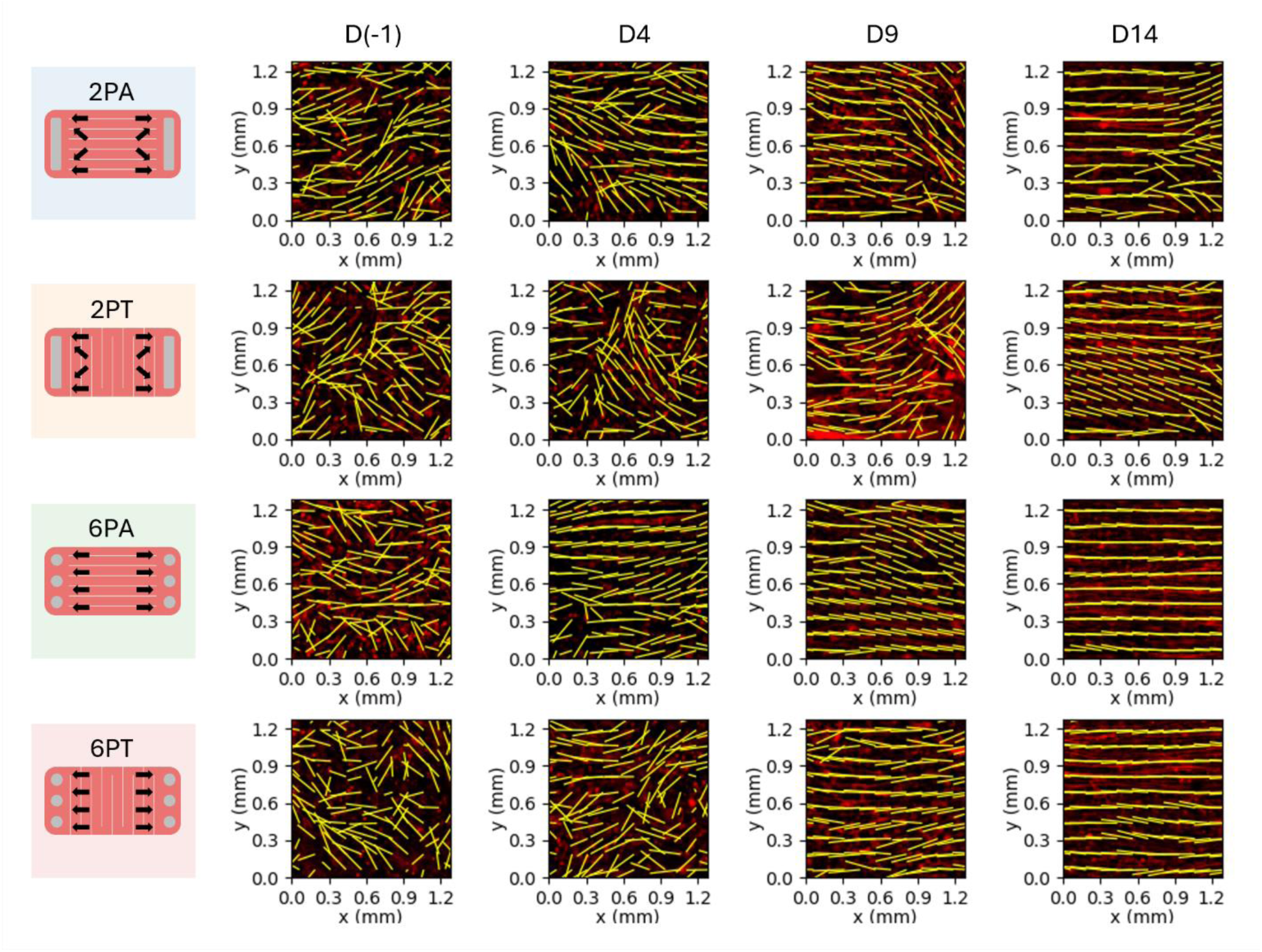
Fibre orientation and alignment over time for different anchoring systems and print path directions. Vector fields of F-actin orientation on top of fluorescence images (10X) of F-actin fibres (red; 5% of the tissue surface represented).

**Fig. S3.**
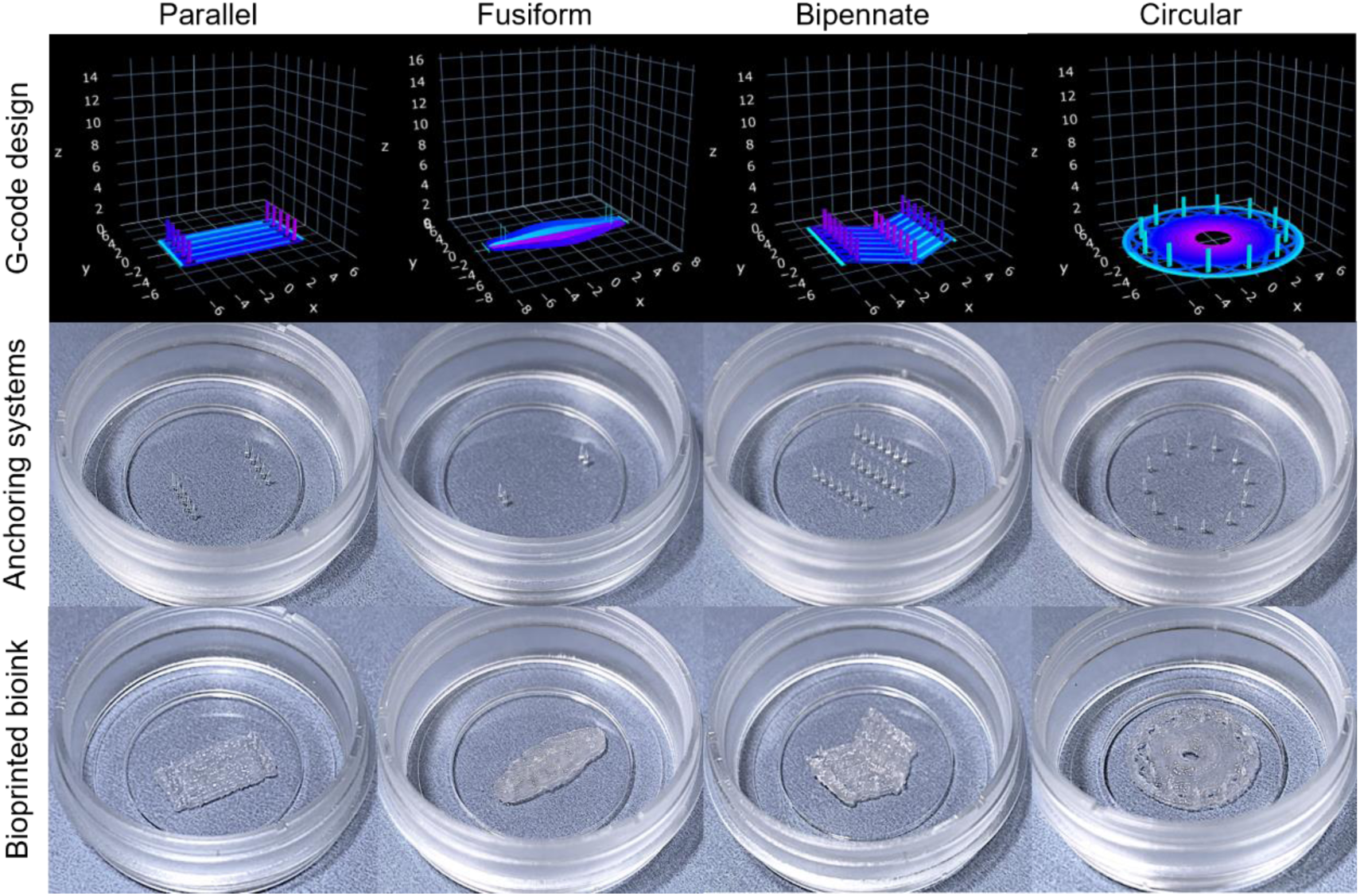
Bioinspired eSMT design and bioprinting. Digital print path, 3D printed PDMS anchoring systems, and bioprinted cell-laden bioink around previously cured pillars. Tissue thickness: 800 µm.

**Fig. S4.**
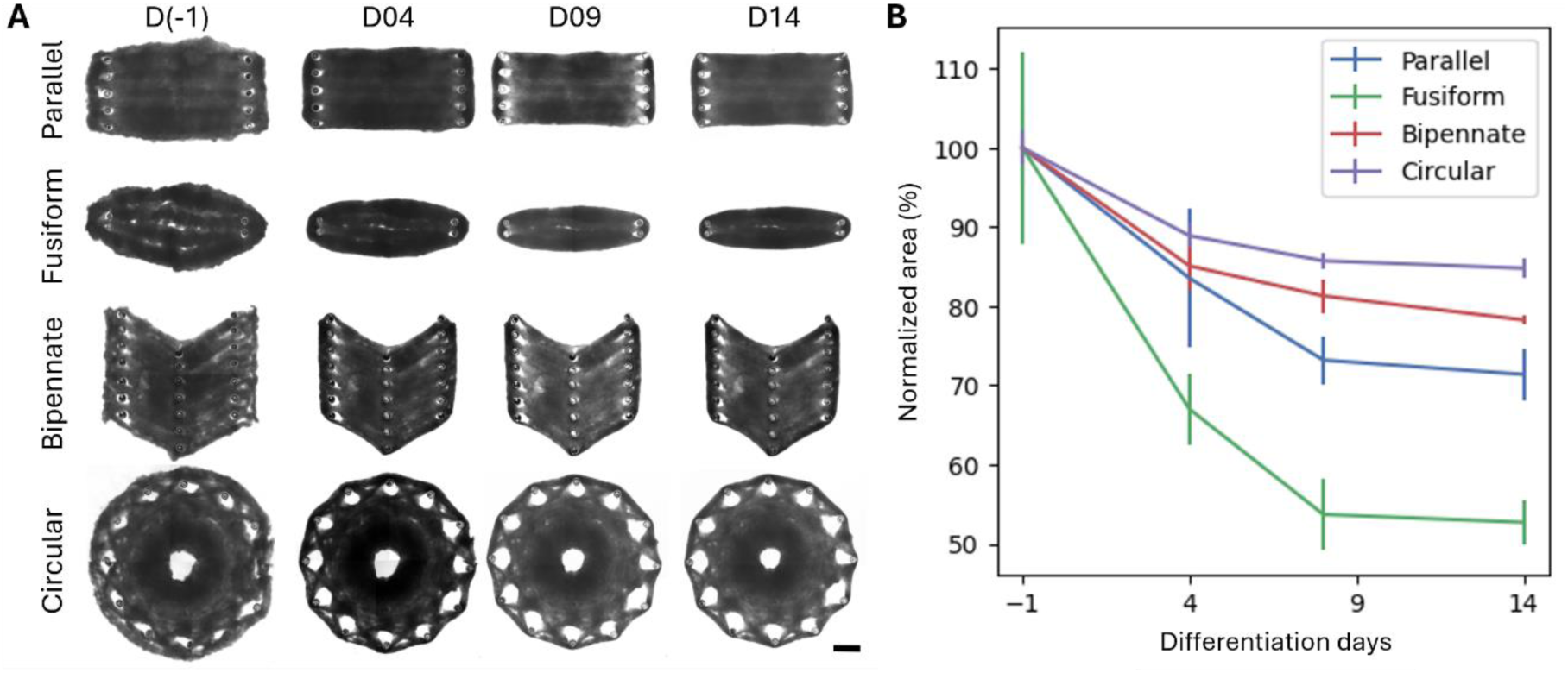
Bioinspired eSMT shape evolution. (**A**) Bright field images (1.25X) of the eSMTs over time. Scale bar: 2mm. (**B**) Evolution of the normalized area of the tissue (area/initial_area) over time (n = 3).

**Fig. S5.**
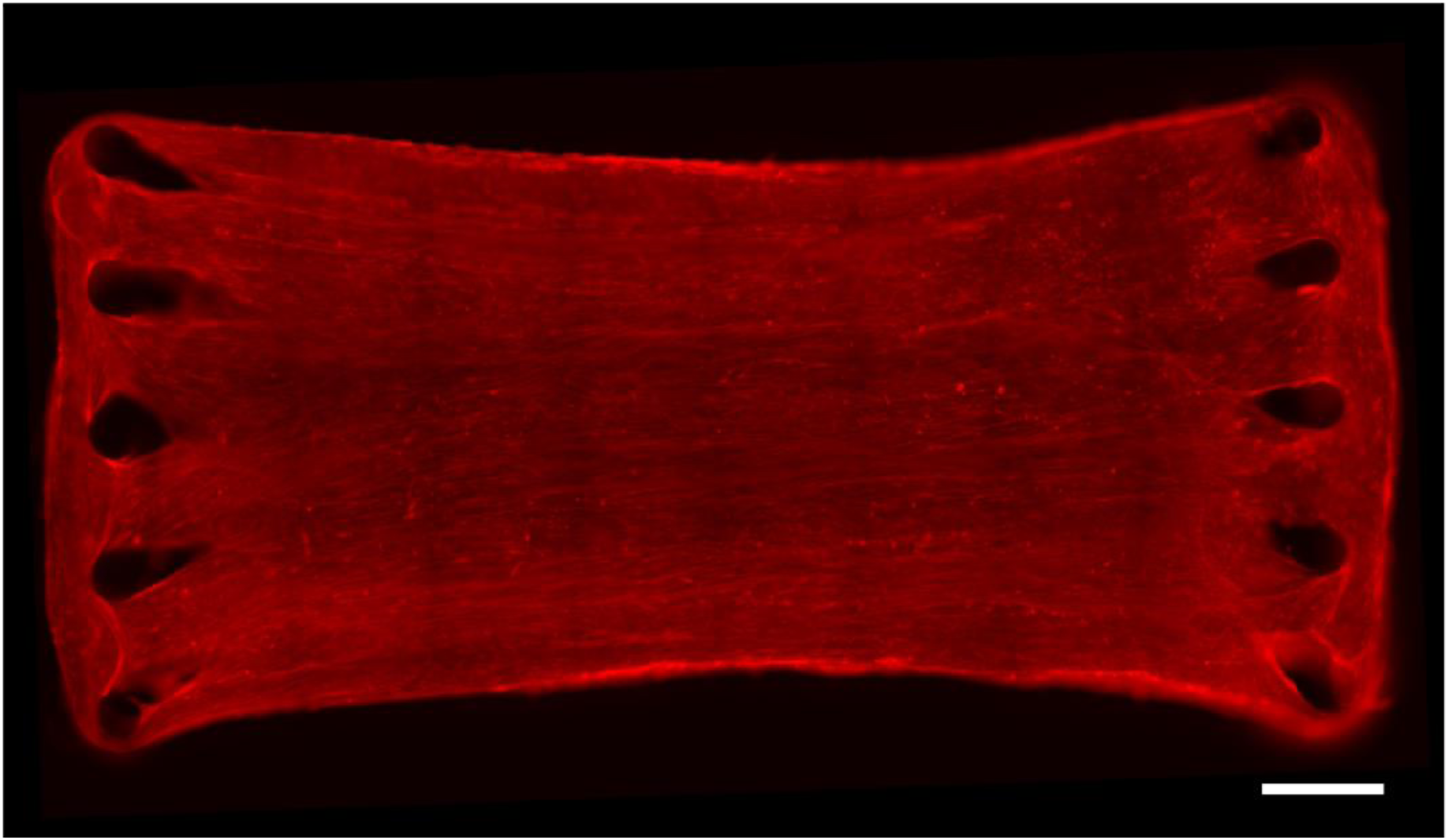
Parallel eSMT fibres. Fluorescence image (10X) of stained F-actin fibres (red). Scale bar: 1mm.

**Fig. S6.**
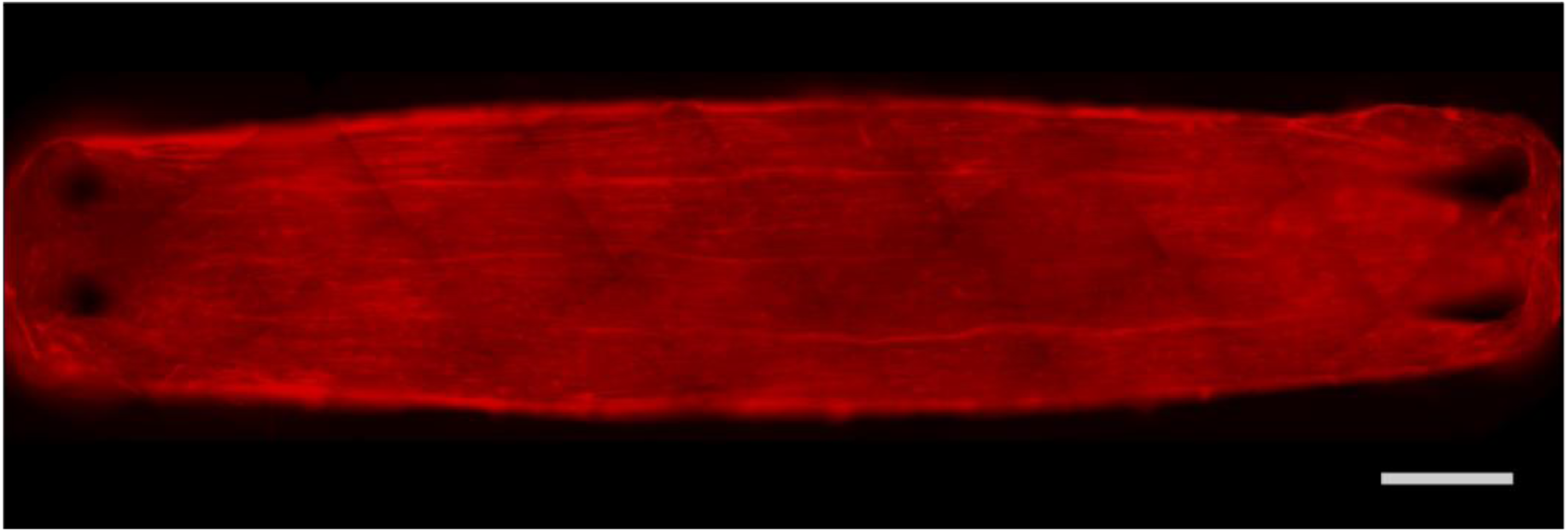
Fusiform eSMT fibres. Fluorescence image (10X) of stained F-actin fibres (red). Scale bar: 1mm.

**Fig. S7.**
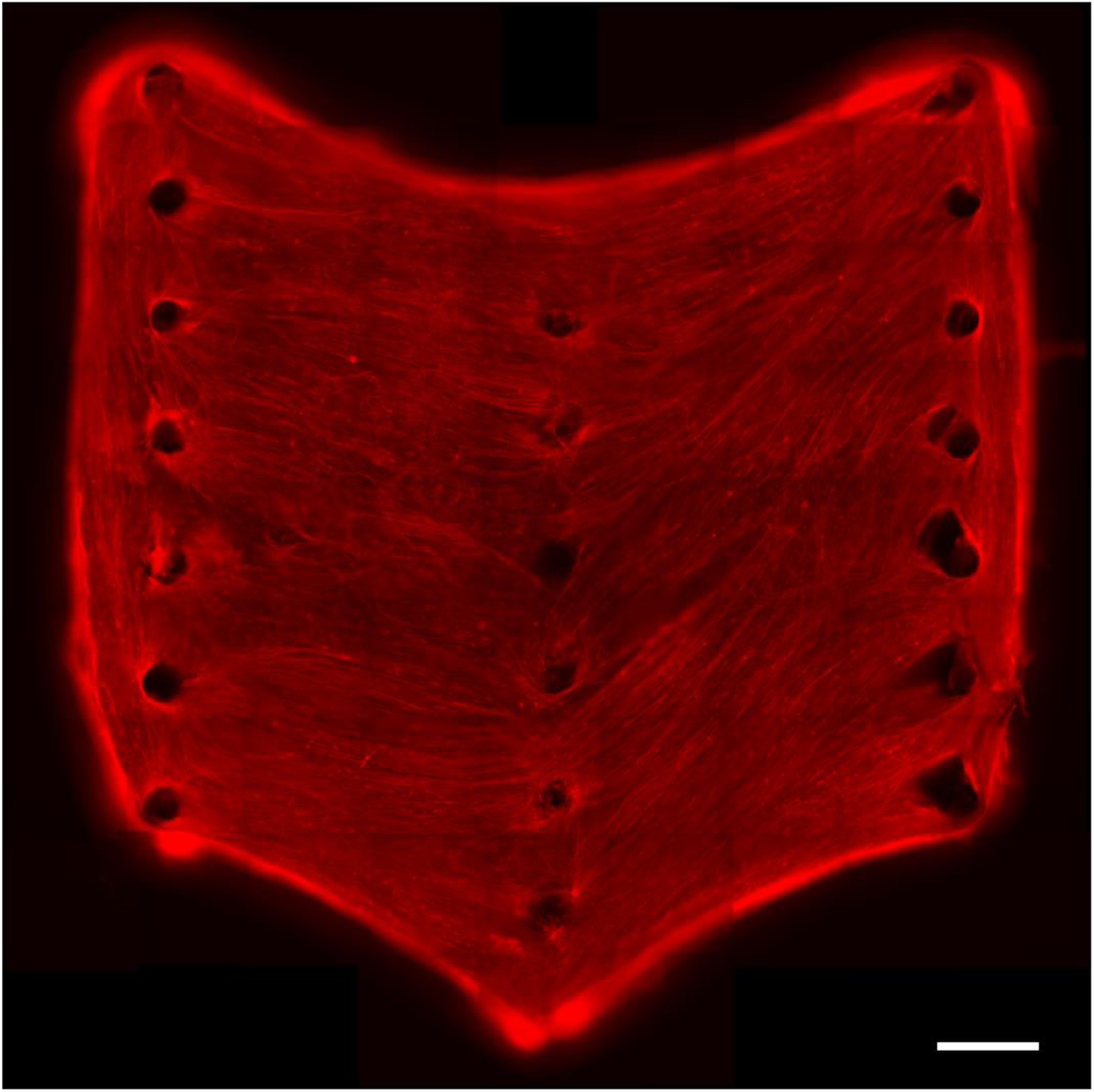
Bipennate eSMT fibres. Fluorescence image (10X) of stained F-actin fibres (red). Scale bar: 1mm.

**Fig. S8.**
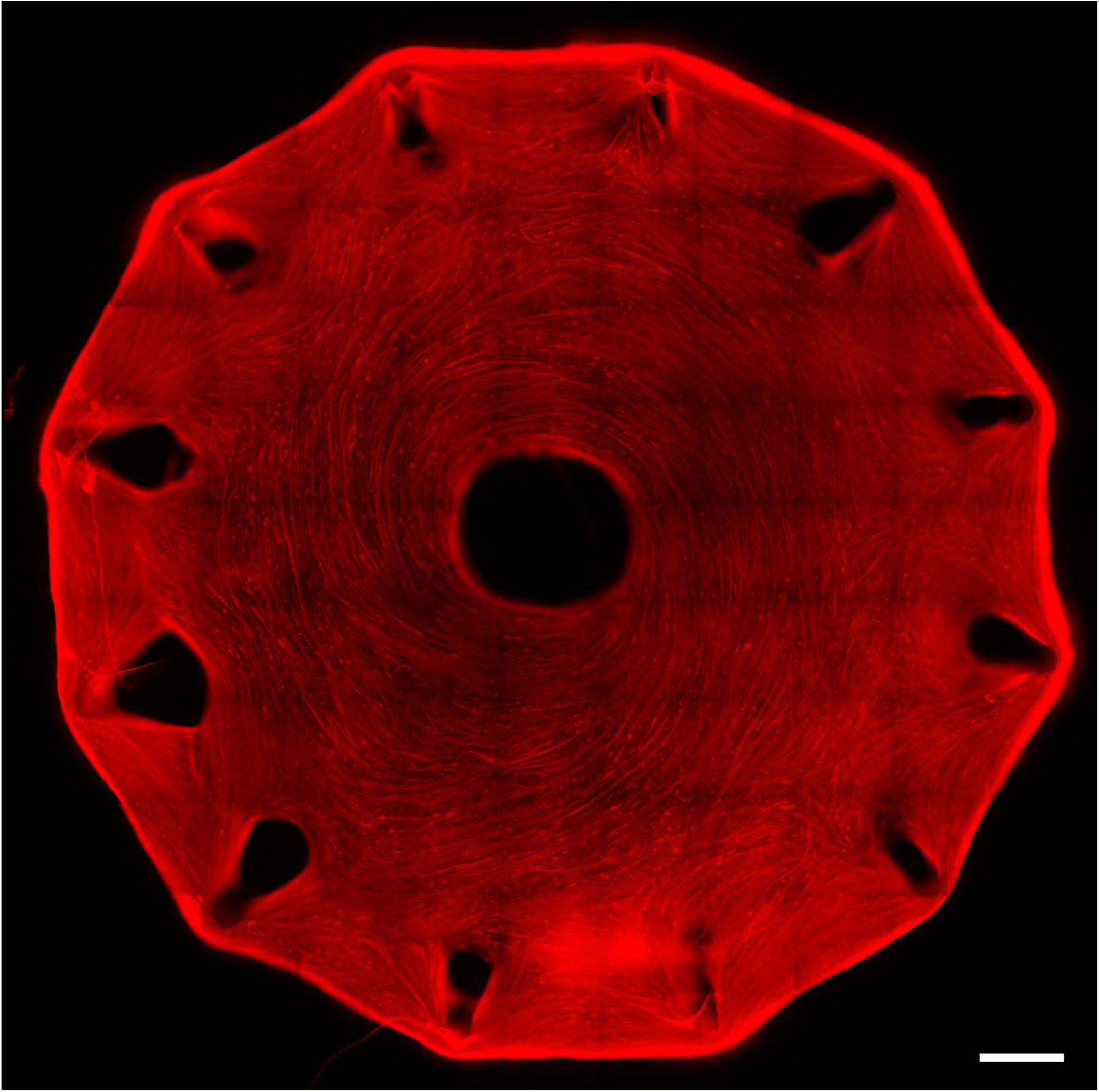
Circular eSMT fibres. Fluorescence image (10X) of stained F-actin fibres (red). Scale bar: 1mm.

**Fig. S9.**
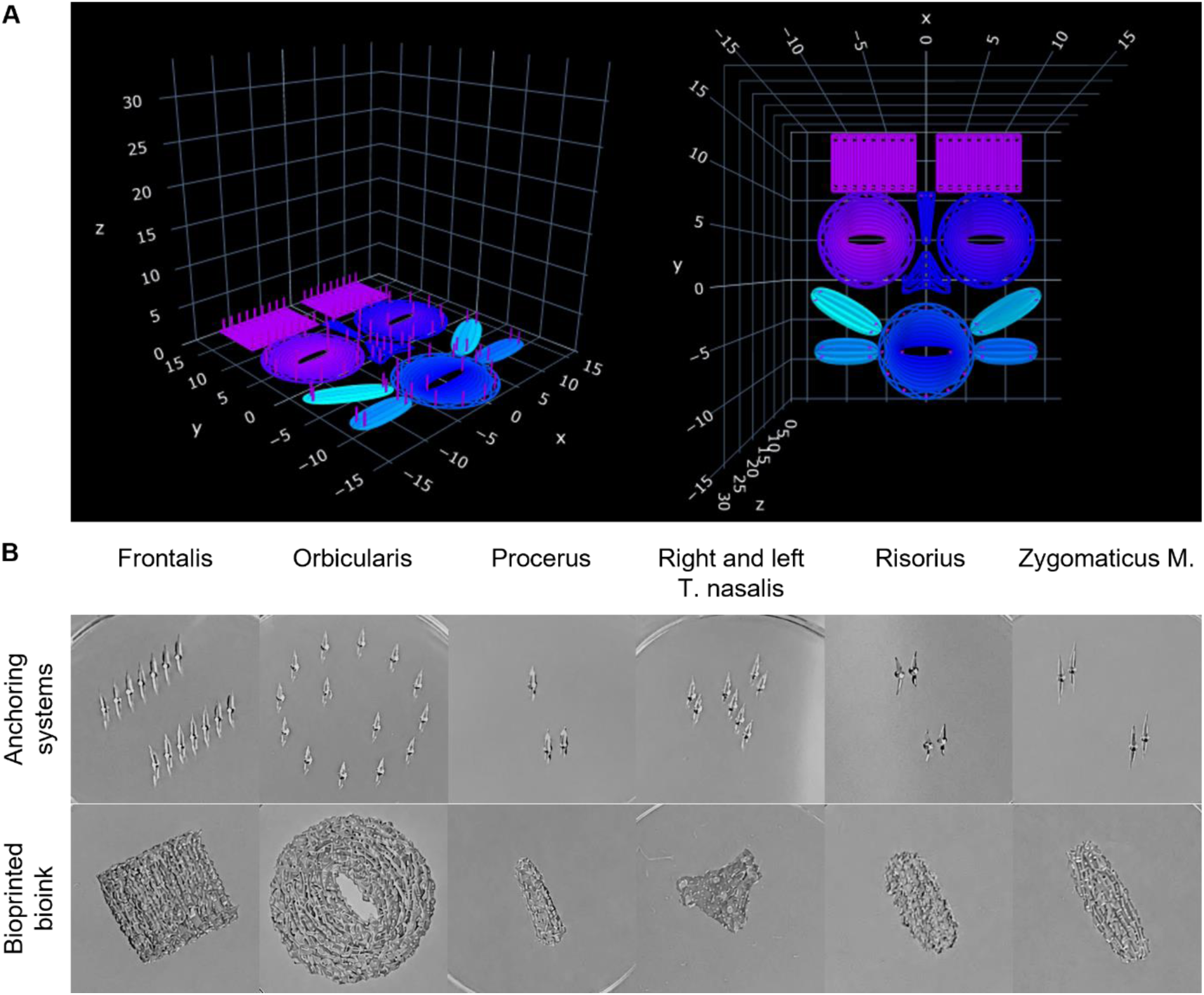
Face musculature design and 3D printing. (**A**) Representations of the digital print paths for pillars and eSMT defined by G-code. (**B**) Pictures of the printed PDMS pillars and bioprinted bioink immediately after printing.

**Fig. S10.**
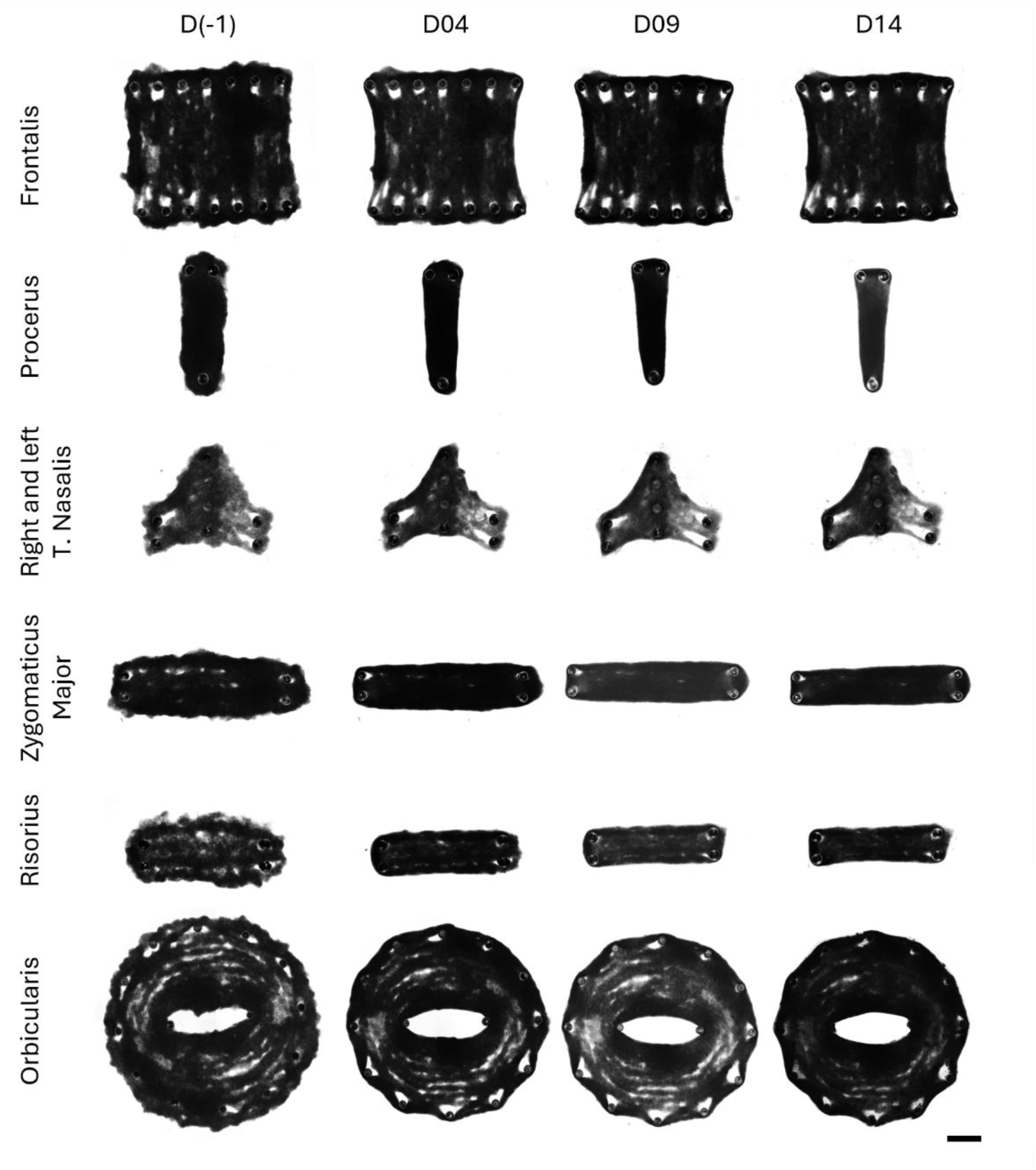
Face eSMTs shape evolution. Bright field images (1.25X) of the facial eSMTs over time. Scale bar: 2mm.

**Fig. S11.**
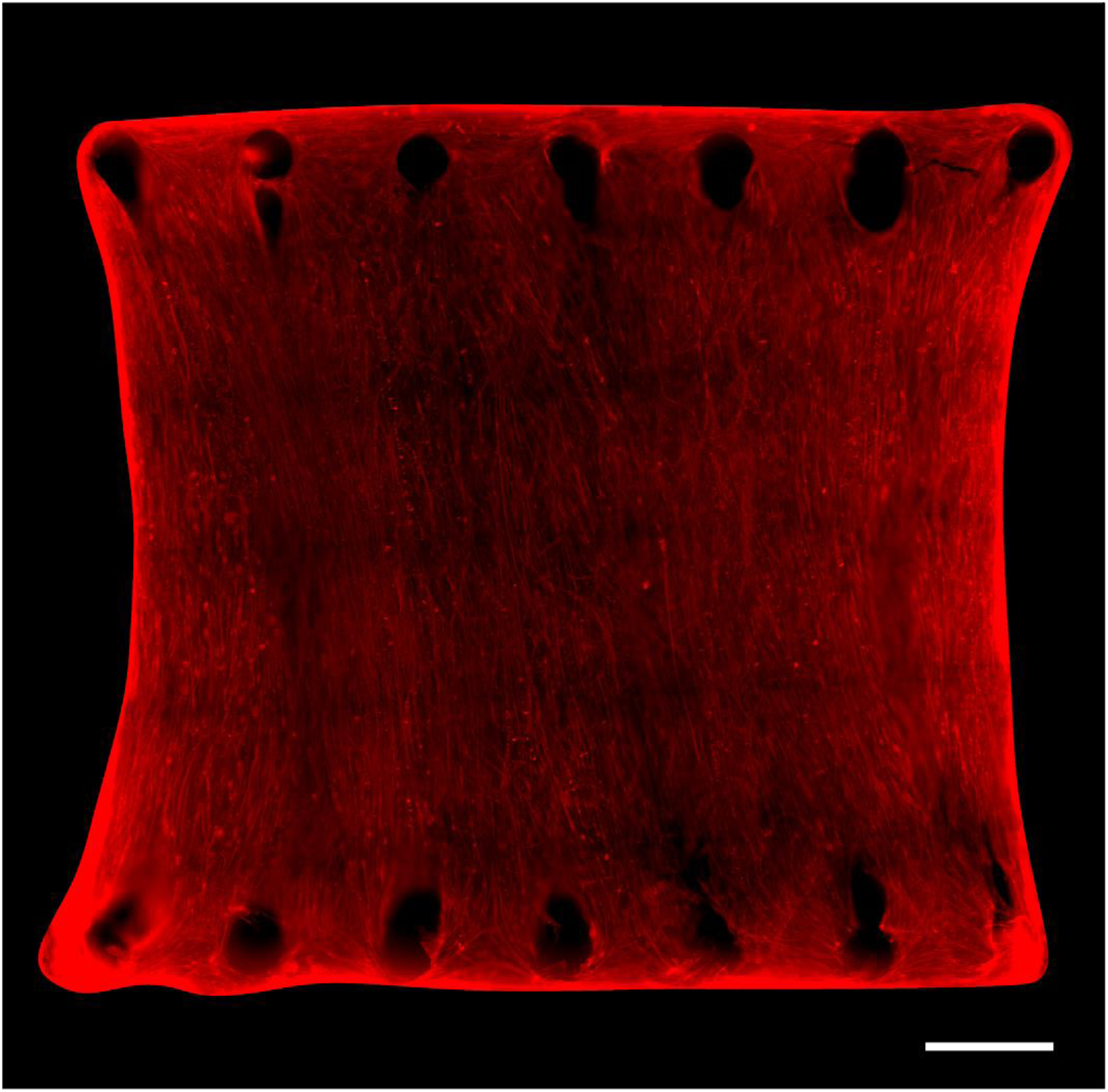
Frontalis eSMT fibres. Fluorescence image (10X) of stained F-actin fibres (red). Scale bar: 1mm.

**Fig. S12.**
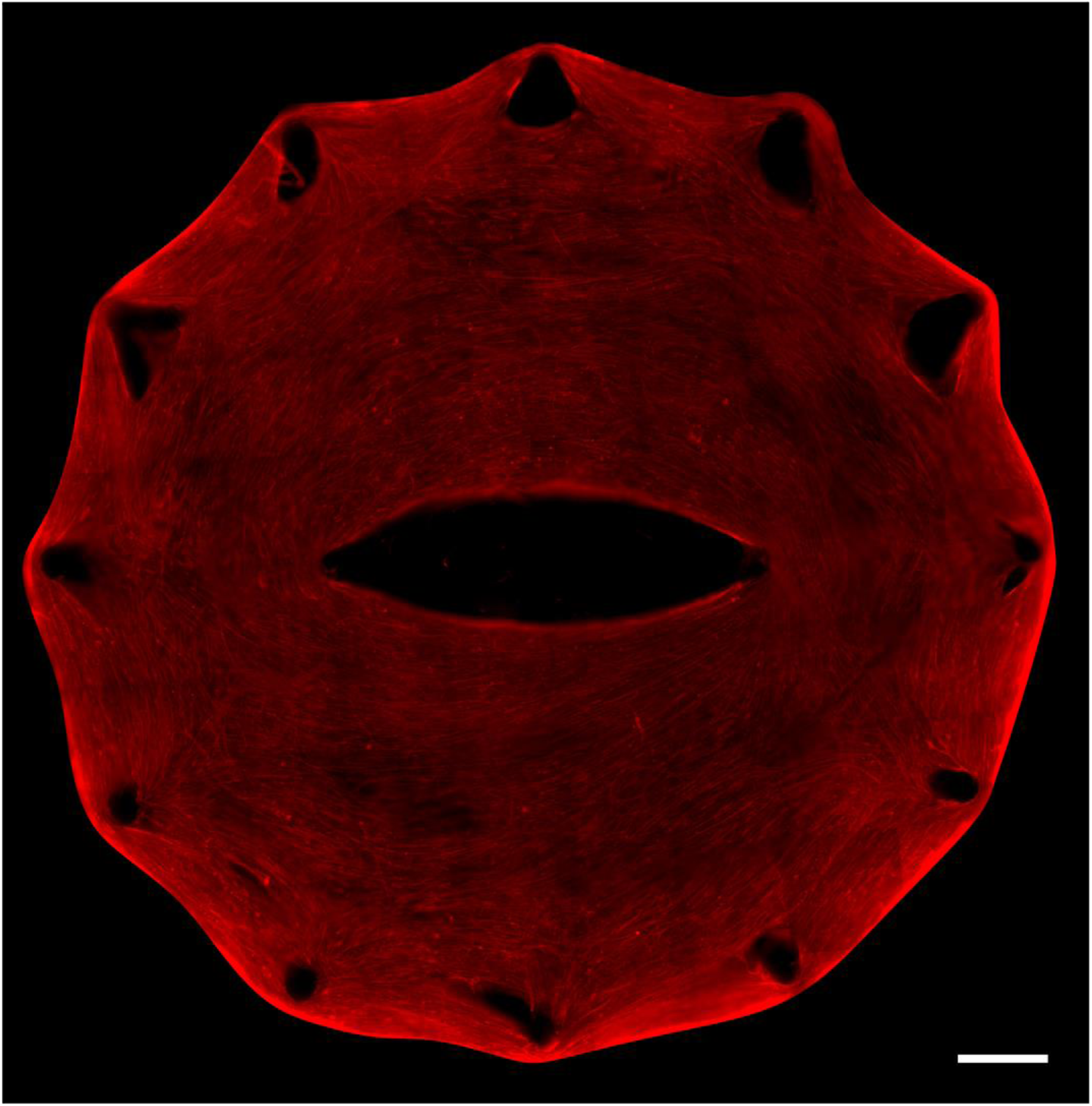
Orbicularis eSMT fibres. Fluorescence image (10X) of stained F-actin fibres (red). Scale bar: 1mm.

**Fig. S13.**
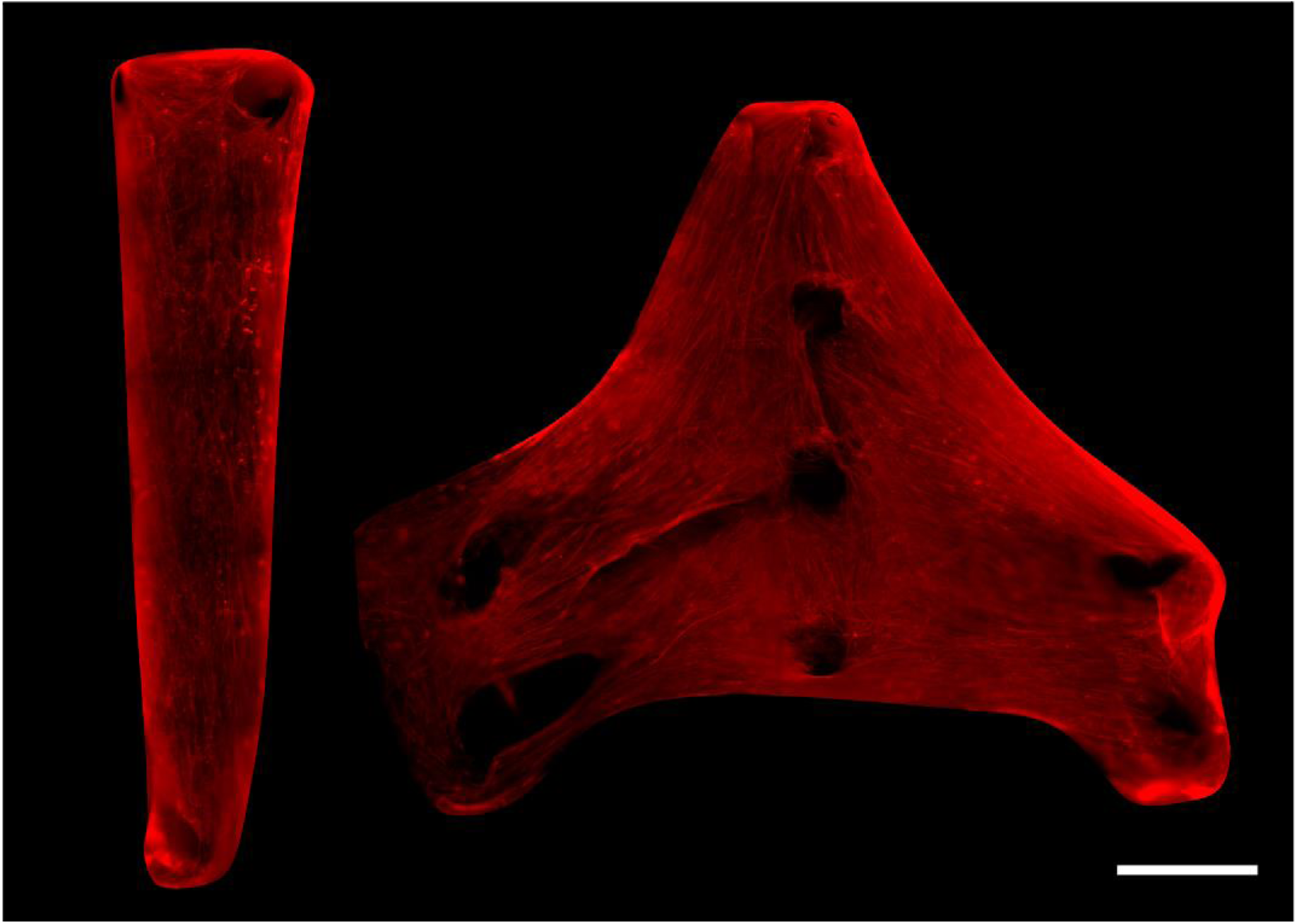
Procerus, and Transverse nasalis eSMT fibres. Fluorescence image (10X) of stained F-actin fibres (red). Procerus eSMT (left), and left and right Transverse nasalis (right). Scale bar: 1mm.

**Fig. S14.**
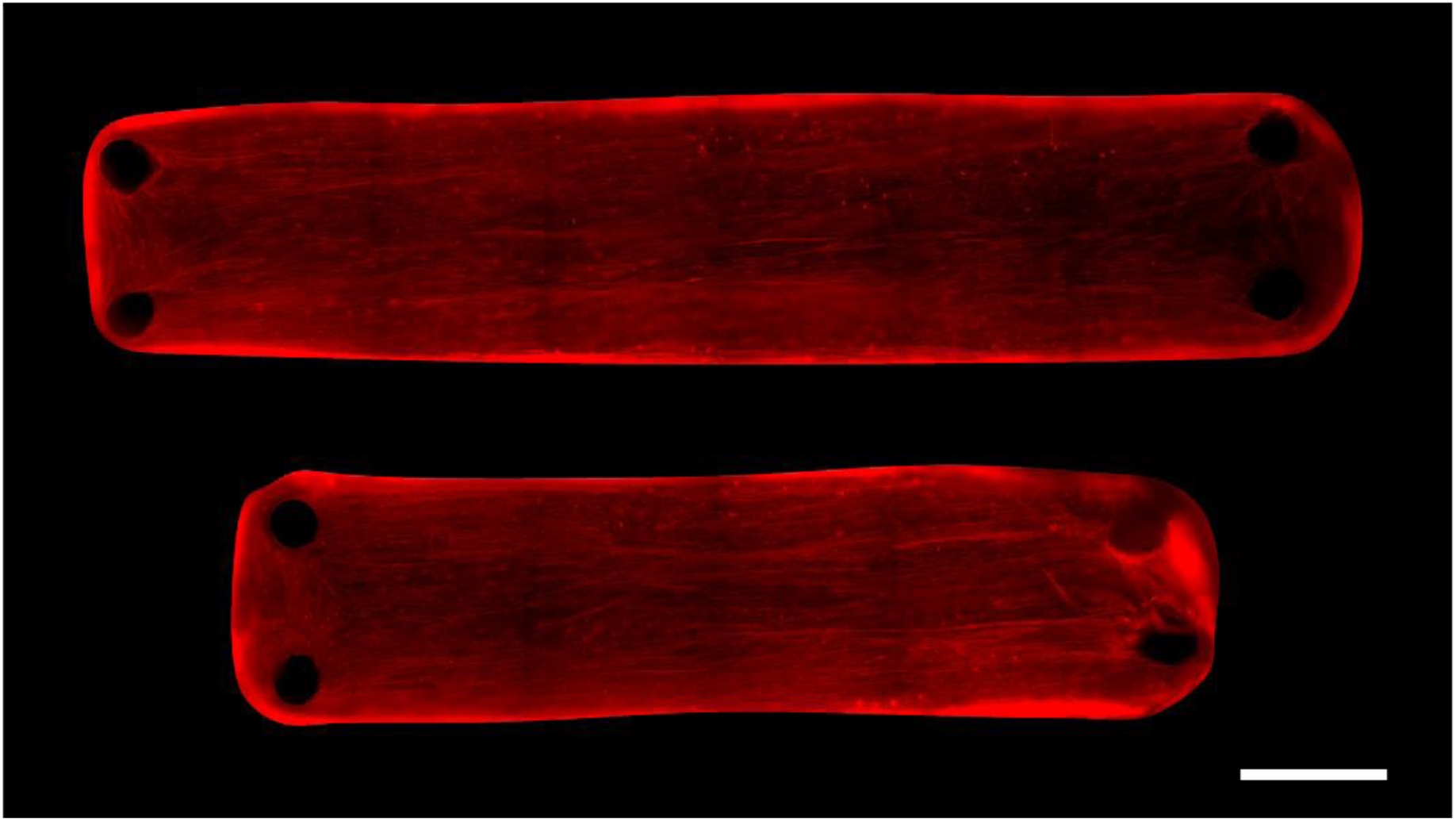
Zygomaticus major and risorius eSMT fibres. Fluorescence image (10X) of stained F-actin fibres (red). Zygomaticus major eSMT (top) and risorius eSMT (bottom). Scale bar: 1mm.

**Fig. S15.**
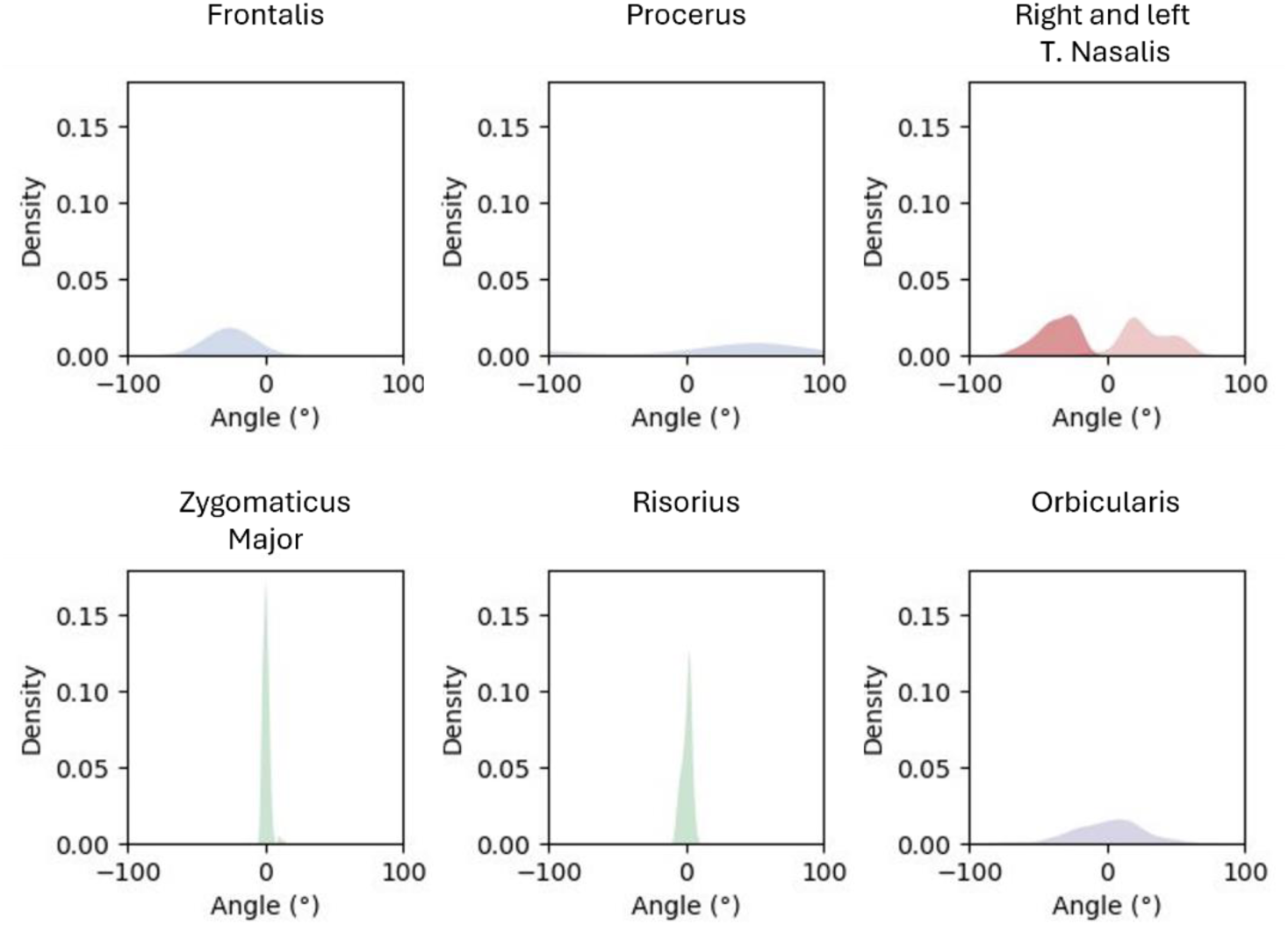
Face eSMTs fibre orientation. Kernel density plots of fibre angles, where 0° corresponds to the horizontal direction in Figures S9-S13, except for the orbicularis design, where 0° represents the direction perpendicular to the radius of the tissue (n = 1).

**Fig. S16.**
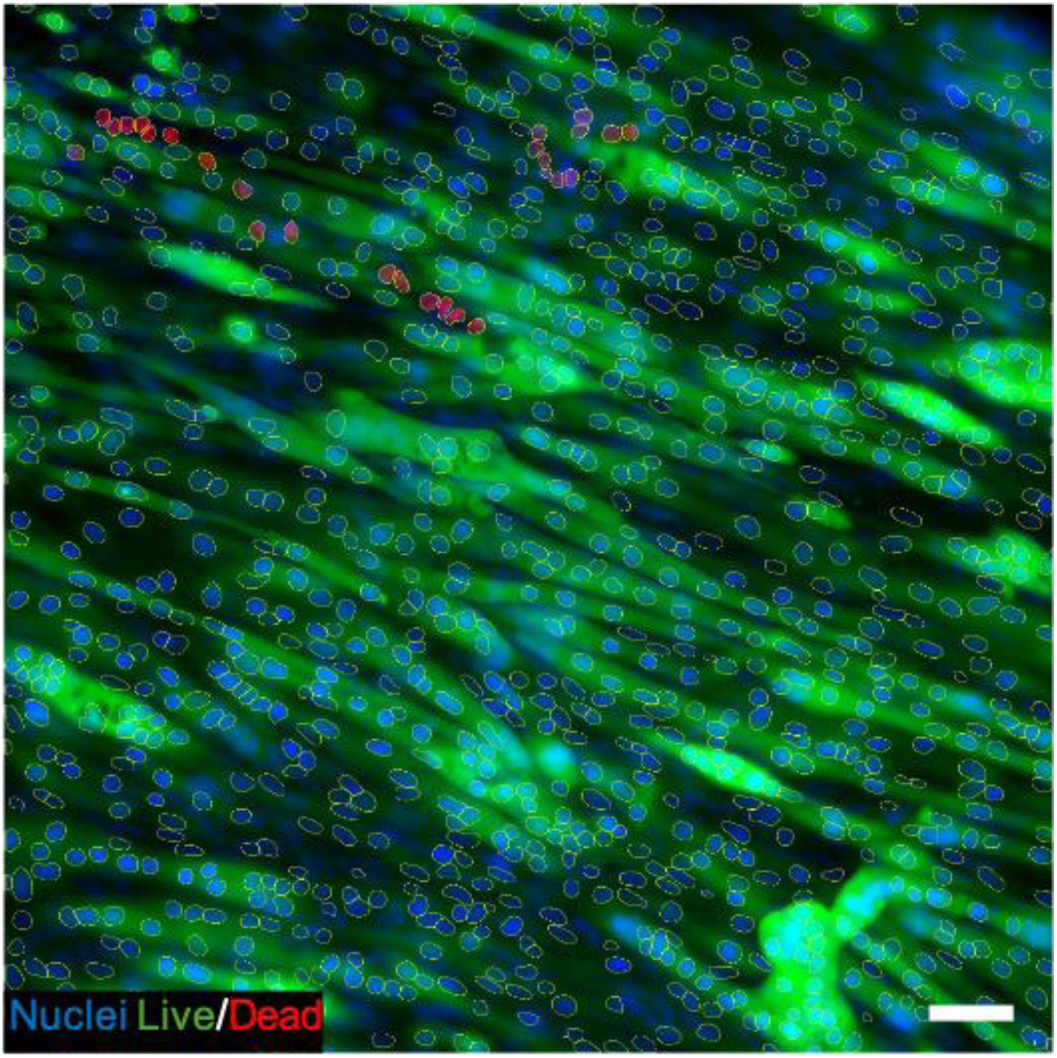
Live/Dead image segmentation. Representative fluorescence image (20X) showing the live/dead staining used to calculate cell viability. Live myofibres are shown in green, the nuclei of dead cells in red, and the rest of the nuclei in blue. Nuclei segmentation is shown in yellow. Scale bar: 50µm.

**Fig. S17.**
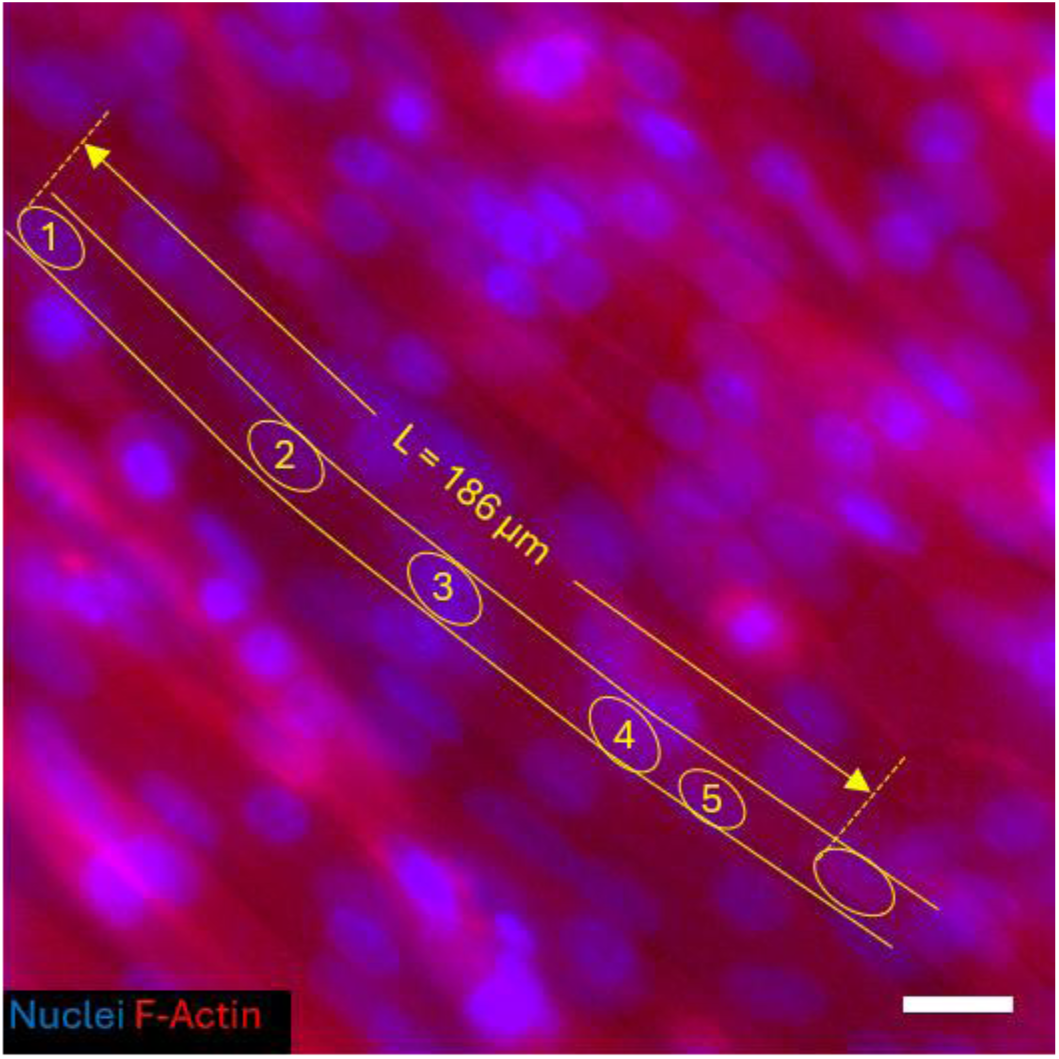
Nuclei density measurement. Representative fluorescence image (20X) used to calculate nuclei density, showing F-actin fibres (red) and nuclei (blue). Nuclei numbered 1 to 5 are consecutive nuclei within a single fibre, and L is the length of the fibre segment containing these five nuclei. For this fibre segment, the nuclei density is therefore 5/L. Scale bar: 20µm.

**Movie S1.** Electrically induced contraction of bioinspired 3D-bioprinted eSMTs. Bright-field videos (1.25X, 25 fps) of twitch contractions of parallel, fusiform, bipennate, and circular tissues at 1 Hz. Scale bar: 1 mm.

**Movie S2.** Electrically induced contraction of 3D-bioprinted facial musculature. Single tetanic contraction and relaxation of self-standing, bioprinted facial muscles after 14 days of differentiation. Scale bar: 2 mm.

**Movie S3.** Electrically induced contraction of 3D-bioprinted facial musculature on 3D-printed skull. Single tetanic contraction and relaxation of bioprinted facial muscles anchored on top of a 3D-printed skull after 14 days of differentiation. Scale bar: 2 mm.

**Movie S4.** Electrically induced contraction of 3D-bioprinted facial musculature on 3D-printed skull: Frontalis contraction. Eyebrow lift animated schematics and single tetanic contraction and relaxation of bioprinted right Frontalis eSMT partially anchored on a 3D-printed skull after 14 days of differentiation Scale bar: 2 mm. The schematic on the left was created with BioRender.com.

**Data S1. (separate file)** Data used to generate Figures 2B-D, corresponding to cell viability, fusion index, and nuclei density over time for different anchoring systems and print path directions.

**Data S2. (separate file)** Data used to generate Figure 3E, corresponding to Cell Viability, Fusion Index, and Nuclei Density after 14 days of differentiation for parallel, fusiform, bipennate, and circular eSMTs.

**Data S3. (separate file)** Data used to generate Figure 4D, corresponding to the contraction of eSMTs with parallel, fusiform, bipennate, and circular architectures.

**Data S4. (separate file)** Data used to generate Figures 2E, 3G, 4F, and S15, corresponding to the local orientation of myofibres and the local orientation of the direction of contraction.

